# Ecosystem Links: Macrophytes, Snail Preferences, and Trematode Transmission in Man-Made Water Bodies

**DOI:** 10.1101/2024.02.29.582719

**Authors:** Aspire Mudavanhu, Emilie Goossens, Ruben Schols, Tawanda Manyangadze, Tamuka Nhiwatiwa, Tine Huyse, Luc Brendonck

**Affiliations:** Department of Biological Sciences, Bindura University of Science Education, Bindura, Zimbabwe; Laboratory of Animal Ecology, Global Change and Sustainable Development, KU Leuven, Leuven, Belgium; Department of Biology, Royal Museum for Central Africa, Tervuren, Belgium; Laboratory of Aquatic Biology, KU Leuven Kulak, Kortrijk, Belgium; Department of Geosciences, School of Geosciences, Disaster and Development, Faculty of Science and Engineering, Bindura University of Science Education, Bindura, Zimbabwe; Discipline of Public Health Medicine, College of Health Sciences, University of KwaZulu-Natal, Durban, 4000, South Africa; Department of Fisheries and Ocean Sciences, School of Agriculture and Fisheries, University of Namibia, Henties Bay, Namibia; Water Research Group, Unit for Environmental Sciences and Management, North-West University, South Africa

## Abstract

Freshwater snails act as obligate intermediate hosts for trematode parasites that cause trematodiases threatening public and veterinary health, and biodiversity conservation. Therefore, interest has re-emerged in snails as a target for disease control, yet their ecology is poorly understood. We studied the relationship between physical and chemical water parameters, macroinvertebrates, macrophytes, land use, and snail abundance, diversity, and infection rate in man-made reservoirs in eastern Zimbabwe. We observed no significant relationship between water quality parameters or macroinvertebrates and snail communities, but a significant association existed between specific macrophytes and snail species. Schistosome-competent snails (i.e., *Biomphalaria pfeifferi* and bulinids) were most associated with emergent *Cladium mariscus,* whereas *Physella acuta* was associated with submerged oxygen weed, *Lagarosiphon major*. This offers a possibility to incorporate the management of macrophytes in integrated snail control schemes. Diversity of freshwater snail species significantly varied across land use types with the lowest observed diversity in the commercial tobacco farm section, dominated by invasive exotic *P. acuta* and *Pseudosuccinea columella*, as compared to the less impacted conserved area, reflecting the adverse effects of agriculture on biodiversity. Out of the 547 schistosome host snails, 88 were shedding cercariae (16.1%) of various types, including schistosomes and amphistomes. We did not find any significant associations between macroinvertebrate or macrophyte diversity and snails and their infection rate.

## Introduction

Snail-borne trematode infections are neglected tropical diseases prevalent in Sub-Saharan Africa (Gyapong & Boatin, 2016). Schistosomiasis is Africa’s most common trematodiasis of public and veterinary health significance. The disease continues to exert pressure against social and economic development, particularly in sub-Saharan Africa where more than 80% of global schistosomiasis cases are concentrated (Hotez et al., 2014; Lai et al., 2015; Steinmann et al., 2006; WHO, 2020). There has been significant advancement in managing schistosomiasis morbidity in Africa, facilitated through large-scale administration of praziquantel to school-aged children and other high-risk groups (Kokaliaris et al., 2022). The antischistosomal drug is only effective in treating mature infections and does not prevent reinfection (Pica-Mattoccia & Cioli, 2004; Sokolow et al., 2017). As such, the disease continually persists, expands, and intensifies, particularly in regions where water resources have been developed, such as large dams and irrigation systems (Gyasi et al., 2019; Steinmann et al., 2006). It was therefore recommended that mass chemotherapy should be complemented with additional efforts, such as sustainable vector control that is adapted to the local environment (Sokolow et al., 2016).

As heteroxenic parasites, trematodes, such as schistosomes that cause schistosomiasis, require freshwater snails as intermediate hosts for the development of free-swimming cercariae, the infective larval stage (Toledo & Fried, 2014; Webbe & el Hak, 1990). Thus, snails are considered pivotal in trematode lifecycles and controlling, primarily through molluscicide but also through environmental modifications, has demonstrated efficiency in reducing the spread of schistosomiasis (Yang et al., 2012) and has resulted in numerous successful control outcomes (King et al., 2015; Rollinson et al., 2013). Chemical-based mollusciciding is expected to play a significant role in the future elimination schemes of schistosomiasis (King et al., 2015; Sokolow et al., 2016). This will be particularly more pronounced in rural areas where most cases of schistosomiasis occur. Additionally, due to the potential zoonotic transmission and the possible role of animals as reservoirs, livestock farms, and wildlife sanctuaries may also be targeted for snail and trematode control. However, most molluscicides used today are not ‘eco-friendly’ as they do not selectively eradicate snails but also harm other aquatic life and persist in the environment (Dai et al., 2008; Keighley et al., 2021; Yang et al., 2020).

Therefore, it is essential to have extensive knowledge of the local ecosystem to evaluate the risks of using chemicals and explore complementary snail control techniques that require minimum yet effective doses. There is a variety of ecological factors influencing snail distribution and infection rate, which in turn may affect disease epidemiology. Studying these ecological factors is of great importance as targeting snails is one of the most impactful measures for disease control (Sokolow et al., 2018). Local conditions such as temperature, water quality, macrophyte cover, and various biodiversity components can directly impact vector snail distribution and life history and may ultimately favor parasite transmission (Becker et al., 2020; Halstead et al., 2018; Kalinda et al., 2017; Malan et al., 2009).

High aquatic community diversity, on the other hand, can locally result in reduced disease risk in several different ways, which is known as the ‘dilution effect’, a hypothesis that has been extensively studied and demonstrated in parasitic diseases (Civitello et al., 2015; Keesing et al., 2006). The presence of non-host species can affect transmission via interactions with either the parasite or the host. For the parasites, these include physical interference with movement, the infection of decoy- or dead-end hosts, and predation on parasite larval stages by invertebrate predators (Hopkins et al., 2013; Johnson et al., 2009). The snail hosts can be affected by competition and predation by other invertebrates (Früh et al., 2017; Larson & Ross Black, 2016; Maldonado & Martín, 2019; Turner et al., 2007).

Ecological disturbances caused by both natural and human factors can lead to the emergence and spread of diseases (Lawler et al., 2021). Activities such as deforestation, human settlement, road construction, and water development projects have been linked to a rise in parasitic diseases (Ellwanger et al., 2020; Johansen et al., 2023; Kibret, 2018; Sokolow et al., 2017). The mechanisms by which these factors increase disease transmission are diverse and can include changes in water quality due to pollution or disruption, alterations in aquatic community structure, or the introduction of new species, and the expansion of snail habitats caused by the proliferation of macrophytes (Carolus et al., 2019; Lange et al., 2013; Steinmann et al., 2006).

Understanding the intricate relationships between snail diversity, infection rates with trematode parasites, and their ecological context remains a critical yet understudied area in freshwater ecology. Most of the information to date comes from controlled laboratory conditions. The research questions driving this investigation revolve around deciphering the impact of land use patterns, water quality, diversity of aquatic plants, and macroinvertebrates in man-made dams on snail diversity and infection rates. Despite significant progress in understanding host-parasite dynamics, gaps persist regarding the specific influences of ecological variables such as land use practices and ecosystem health on the prevalence and diversity of trematode infections in natural snail populations.

This study aims to unravel the complex interplay among snail communities, infection rates with trematode parasites, and multifaceted environmental factors within man-made reservoirs in a study region of Zimbabwe. The study seeks to bridge the research gaps by exploring the nuanced associations between environmental factors, snail communities, and trematode infections, contributing crucial insights into disease dynamics and ecosystem health within man-made reservoirs. We hypothesized that man-made reservoirs with poor water quality would provide favorable conditions for the snail-parasite complex to thrive, and that macrophytes would be an important predictor of snail diversity. We expected that land use types with human disturbance such as agriculture and settlements would alter the ecological state of the reservoirs, through fertilizer and herbicides regulating snail density through bottom-up effects by increasing algal densities which snails feed upon (Halstead et al., 2018).

## 2. Methods

### 2.1 Study area

Our study was carried out at Imire Rhino and Wildlife Conservancy and the neighboring villages of Mhakwe (Fig. 1). This area is located in the Wedza district of Zimbabwe, some 70 km from the capital city Harare, characterized by a temperate highland tropical climate that typically receives about 136 mm of annual rainfall and average temperatures of 21.5 °C. This region was selected as it is reported to have the second-highest schistosomiasis prevalence in the country (Midzi et al., 2014), coupled with a history of potentially parasite-linked animal mortalities at the conservancy, which presented possibilities to monitor and study snail-borne parasites in relation to land use and environmental conditions. The conservancy is divided into four parts that are characterized by different land use types and wildlife populations (Fig. 1). There are three fenced parts in the conservancy where the main activities are nature conservation and tourism: *herein* referred to as Chiwawe, the Middle section, and Lodge section. The fourth part is a commercial tobacco farm managed by the conservancy and with no wildlife. Three more sites were sampled outside Imire in the surrounding villages, predominantly used for cattle drinking, fishing, irrigation, and household activities. A total of 19 sites were sampled after the warm wet season, in March 2022. All sampled sites are man-made reservoirs consisting of earth dams and concrete structures.

**Fig. 1.**
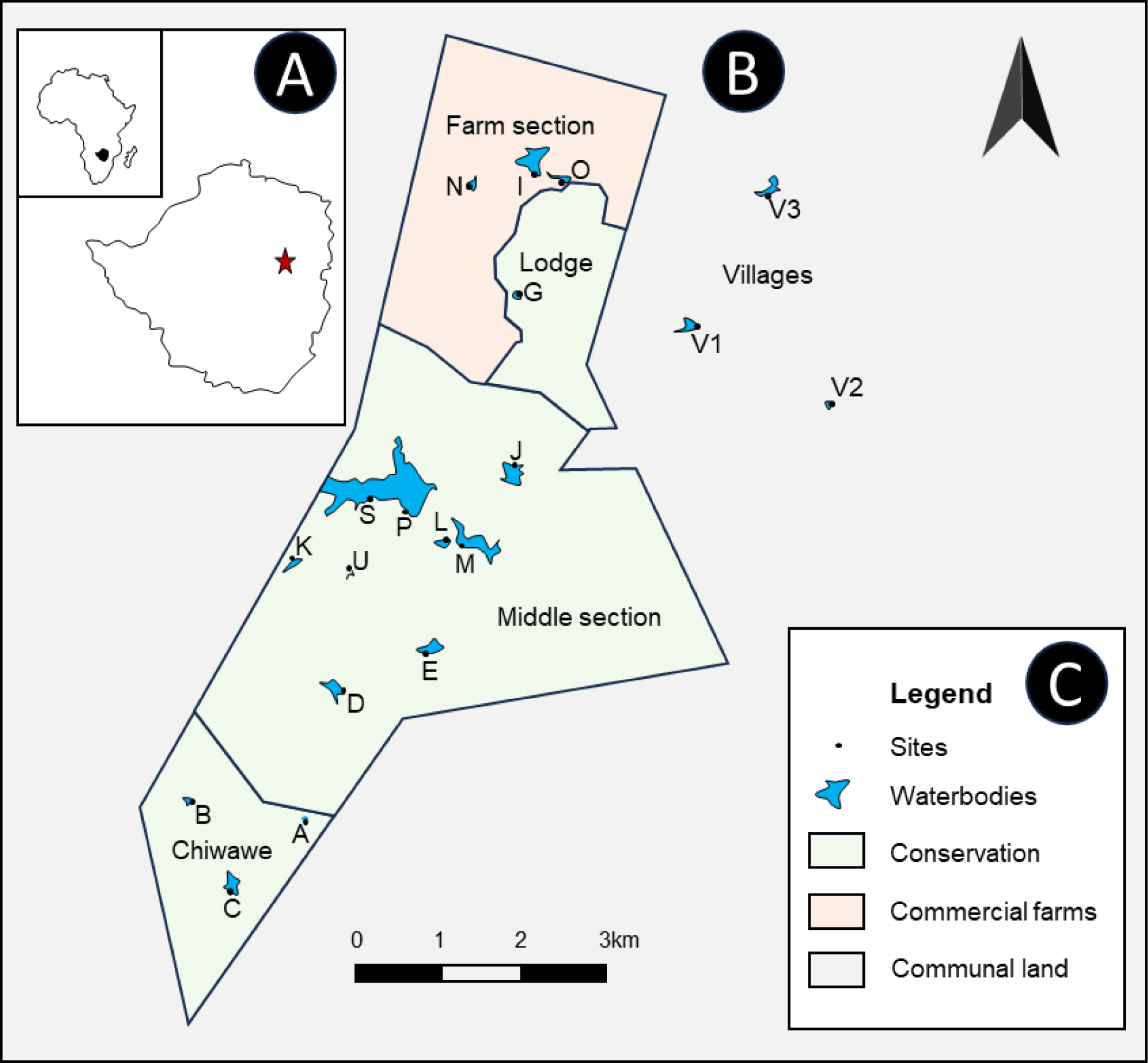
Map of the study area. (a) position of the study area in the southern eastern country of Zimbabwe on the African map (shaded black) and the location of the study region, east on the Zimbabwean map (shown as a red star). (b) Map of Imire showing the sampled sites. The borders of Imire and the different sections within are indicated by black lines. The five parts of the study area are indicated namely Chiwawe, Middle section, Lodge section, Farm section, and the Villages. A scale bar is given to show the distance between sampling points and the arrow shows the direction of North. (c) Legend provides clarity for the Imire map. A total of 19 sampling sites within and outside the confinements of Imire are shown on their respective man-made reservoir of variable sizes (shown in blue color). Land use for each subsection in and around Imire is given as color codes with green representing the area under conservation, peach representing the commercial farm section and grey around Imire representing the communal area.

### 2.2 Snail community structure

Snails were collected by handpicking (wearing gloves) and with 1-mm mesh size scooping nets microhabitats as guided in a standardized way. Collected snails were sorted and further identified based on the diagnostic keys (Brown, 1994; Mandahl-Barth, 1962). Our snail specimens were compared with molecularly identified specimens from the precursor study of Mudavanhu et al., (in press) to combat the extensive phenotypic plasticity that often complicates the identification of freshwater snails based on morphology.

### 2.3 Shedding experiment – cercariae collection

Following overnight storage in darkness, snails were exposed to bright light to mimic daylight, a process known to trigger the emergence of cercariae from their intermediate host - the so-called “*shedding experiment*”. Emerging cercariae were identified at the family level according to appropriate keys (Appleton & Miranda, 2015; Frandsen & Christensen, 1984). Shedding snails were stored individually, while individuals that did not release cercariae (non-infected or pre-patent infection) were pooled together per species per site in 80% ethanol. One week later, ethanol was replaced to ensure optimal conservation for later DNA analysis. Cercariae recovered during the shedding experiment were stored in 80% ethanol in barcoded 1.5ml screw-cap tubes and transferred to the University of Leuven (KU Leuven) and the Royal Museum for Central Africa (RMCA) in Belgium for digitalization and molecular identification.

### 2.4 Snail infection rate

DNA was extracted from snail tissue using the E.Z.N.A.® Mollusc DNA Kit (OMEGA Bio-Tek, Norcross, GA, USA) following the manufacturer’s instructions. The elution process yielded 100 μL of DNA extract. Medical and veterinary significant snails were subjected to DNA extraction for infection examination using PCR. Up to 30 specimens of medically significant snails underwent the multiplex rapid diagnostic polymerase chain reaction (RD-PCR) methods described by Schols et al., (2019) and Carolus et al., (2019). These methods enable detection of *Schistosoma* spp. or *Fasciola* spp. infections in snail DNA extracts. The PCR reaction amplifies multiple markers, including an internal control for snail 18S rDNA, a general trematode primer pair, and species-specific primers for *Schistosoma* or *Fasciola*. Positive *Schistosoma* spp. samples underwent species-specific multiplex PCR, differentiating between various schistosomes of medical and veterinary importance (*S. haematobium*, *S. mansoni*, *S. mattheei*, and *S. bovis*/*S. curassoni*/*S. guineensis*) by generating species-specific COI amplicons of different lengths.

### 2.5 Environmental variables

GPS coordinates were recorded at each site using a Garmin GPSMAP® 60CSx. Water physical and chemical variables were collected using a HANNA instrument® HI9829 multiparameter for pH, temperature (°C), conductivity (µS/cm), salinity (PSU), oxygen (% and mg/L), total suspended solids (mg/L) and total dissolved solids (mg/L). A beaker was used to collect an integrated 10-L water sample from different locations at each sampling site to get a representative sample (Table A.1). Multimeter measurements were done in this 10-L sample. A Turner® AquaFluor fluorometer was used to measure turbidity (NTU), chlorophyll A (μg/L) (as a proxy for biomass of micro-algae), and phycocyanin (μg/L) (as a proxy for biomass of cyanobacteria) concentrations. Furthermore, a 500-ml sample from the 10-L bucket was filtered over a 64-µm zooplankton net and analyzed using a spectrophotometer (HACH DR/2010, Colorado) following standard methods HACH (2007) for sulfates (mg/L), total phosphorus (mg/L), reactive phosphorus (mg/L), nitrates (mg/L), nitrites (mg/l), total nitrogen (mg/L), ammonia (mg/L), chemical oxygen demand (mg/L), calcium (mg/L) and magnesium (mg/L) (Table A.1). Finally, the dam dimensions i.e. depth and area were estimated using a calibrated stick and the ArcGIS application respectively.

### 2.6 Macrophyte and macroinvertebrate community structure

Macroinvertebrates were identified based on Gerber et al., (2004) and their relative abundances were recorded. In addition, the percentage of cover by submerged, emergent, and floating vegetation was visually estimated following the guide by Oldham et al. (2000). Macroinvertebrates were collected by sweeping all (micro)habitats (e.g., bottom, submerged vegetation, floating vegetation, water column) with a handheld 250-µm net for 5-10 minutes relative to dam size. After being separated from debris, macroinvertebrates were stored in 80% ethanol and classified per type using diagnostic keys (Day et al., 2003; de Moor et al., 2003a, 2003b; Stals & de Moor, 2007). Their absolute abundances were recorded.

### 2.7 Data analysis

All data were analyzed using RStudio® (Version 2021.09.0 Build 351). The significance threshold for statistical tests was P = 0.05. Graphs generated in R were exported to Microsoft PowerPoint® to improve layout and readability using the *export* package (Wenseleers, 2016). Shannon diversity *H* was calculated for snails, macroinvertebrates, and macrophytes using the function *diversity* in package *vegan* (Oksanen et al., 2022). To explore similarities in snail communities between sites, Nonmetric Multi-Dimensional Scaling (NMDS) and hierarchical cluster analysis were used. NMDS was performed using the *metaMDS* function from the package *vegan* (Oksanen et al., 2022). Cluster analysis was done using the *hclust* function from the package *stats* (R Development Core Team, 2018). Chord’s distance was used as a distance measure and the clustering method was unweighted average linkage agglomerative clustering (UPGMA). The same methods were used for NMDS and cluster analysis of macroinvertebrate and macrophyte communities. To assess the variation among sites in their physical and chemical variables, a principal component analysis (PCA) was performed using the function *pcrcomp* from package *stats* (R Development Core Team, 2018). The function *fviz_pca_biplot* from package *factoextra* (Kassambara & Mundt, 2020) was used to construct the resulting biplot. Finally, redundancy analysis (RDA) was used to test whether physical and chemical variables, macroinvertebrate community, and/or macrophyte community were (significantly) associated with snail community composition. This was done using the function *rda* from package *vegan* (Oksanen et al., 2022). Prior to this, all community datasets were Hellinger transformed. The significance of the RDA was assessed using permutation F-test statistics from *anova.cca* and the resulting triplots were made using the function *triplot* in the package vegan (Oksanen et al., 2022).

To assess what factors, correlate with snail diversity, abundance, and infection rate, (generalized) linear mixed models were used. For the response variable snail diversity, linear mixed models (LMM) were used, while for count data (richness and abundance) and proportional data (infection rate), generalized linear mixed models (GLMM) were used with a Poisson and binomial distribution, respectively. PCR infection results were used for the variable infection rate. Explanatory variables included land use category, water quality variables, richness, and diversity of macroinvertebrates and macrophytes, and macrophyte cover. One random factor including the part of the study area nested in conservation status (protected/unprotected) was added to the models. We thoroughly examined all models to identify overdispersion and incorporated an additional dispersion parameter when necessary. In the case of generalized linear models, we employed a quasipoisson approach for count data and a quasibinomial for infection rate data. For generalized mixed linear models, we addressed overdispersion by running a secondary model that included a random effect at the observation level. The selection of the best model was based on the comparison of AIC values. Linear mixed models were fitted with the function *lmer* and generalized linear mixed models with the function *glmer*, both from package *stats* (R Development Core Team, 2018). Additional Pearson correlation tests were performed to further investigate some relationships using the function *cor.test* from package *stats*.

## 3. Results

### 3.1 Snail variables (diversity, abundance, and spatial distribution)

A total of 926 freshwater snails from 10 species were collected (Fig. 2). Among them, 547 were identified as schistosome-competent species known to transmit schistosomiasis: *Bulinus tropicus*, *Bulinus truncatus*, *Bulinus globosus*, *Bulinus forskalii*, and *Biomphalaria pfeifferi* (Fig. 2: a-e). The remaining snails consisted of *Radix natalensis*, *Gyraulus* sp., *Melanoides tuberculata*, and the exotic invasive species *Pseudosuccinea columella* and *Physella acuta*.

**Fig. 2.**
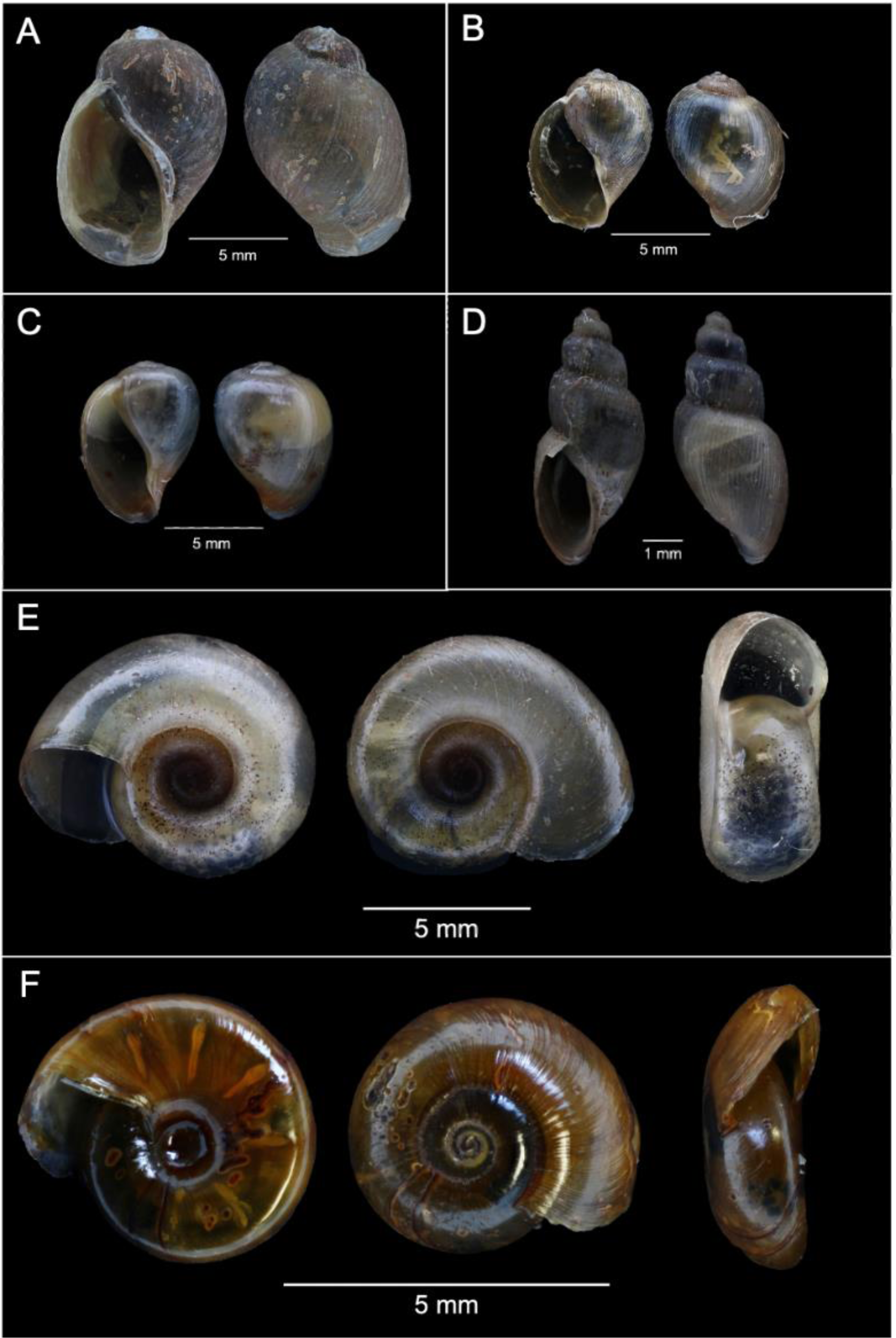
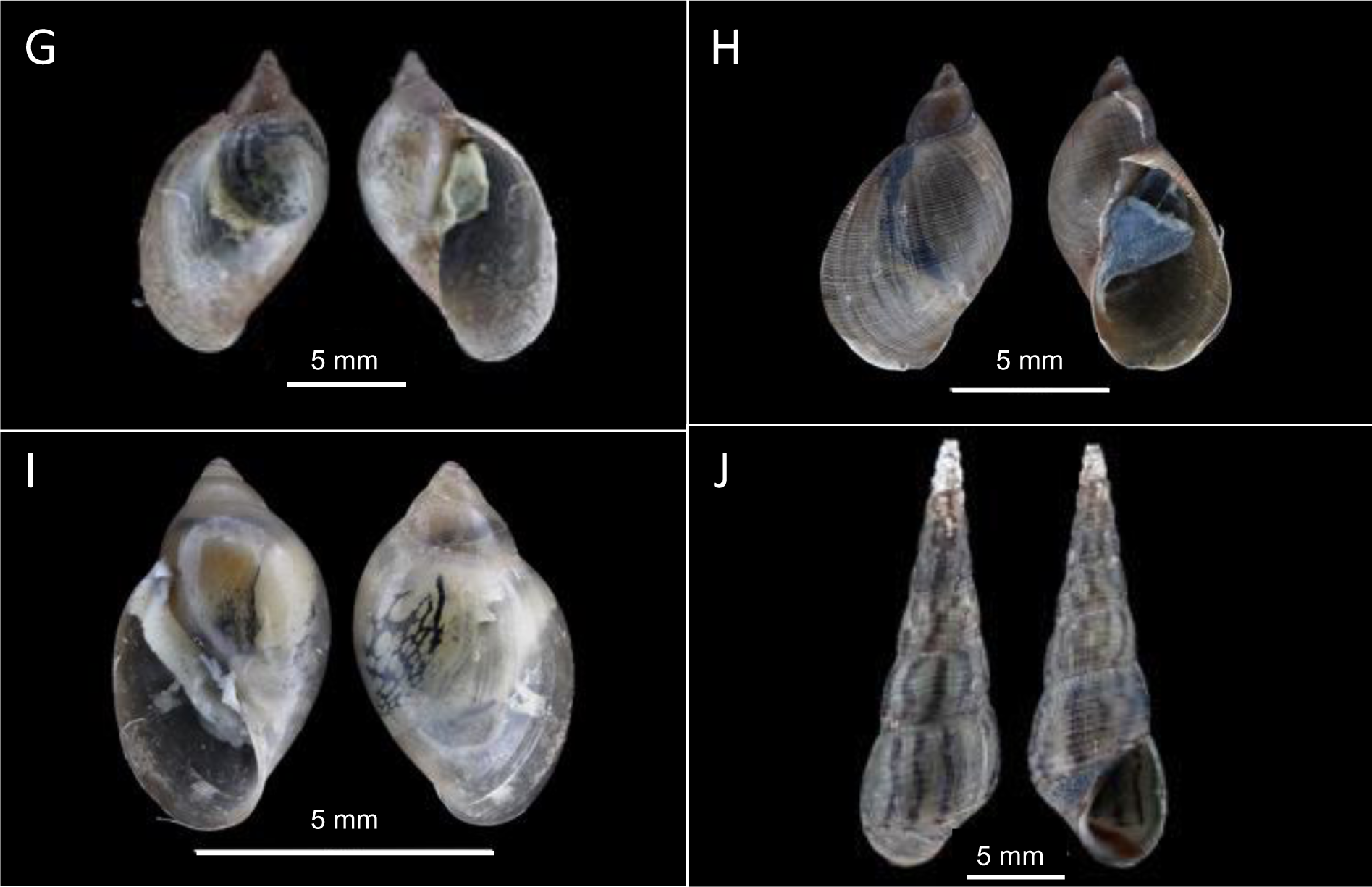
Photographs of snail species collected from man-made reservoirs in Imire. Scale bars indicating either 1mm or 5mm are provided. Species presented are (A) *Bulinus tropicus*, (B) *Bulinus truncatus*, (C) *Bulinus globosus*, (D) *Bulinus forskalii*, (E) *Biomphalaria pfeifferi*, (F) *Gyraulus* sp., (G) *Radix natalensis*, (H) *Pseudosuccinea columella,* (I) *Physa acuta* and (J) *Melanoides tuberculata*.

Snail abundance significantly varied across the five parts of the study area (Χ^2^_(4,*N*=19)_= 14.44, p = 0.006), as illustrated in Fig. 3A. Shannon diversity (H) ranged from 0 to 1.61, with the highest diversity observed in the middle section and villages, and the lowest in the farm section and Chiwawe. Snail abundance ranged from 0 to 173 per site, with the highest abundance in villages and the lowest in the farm section (Fig. 3B).

**Fig. 3.**
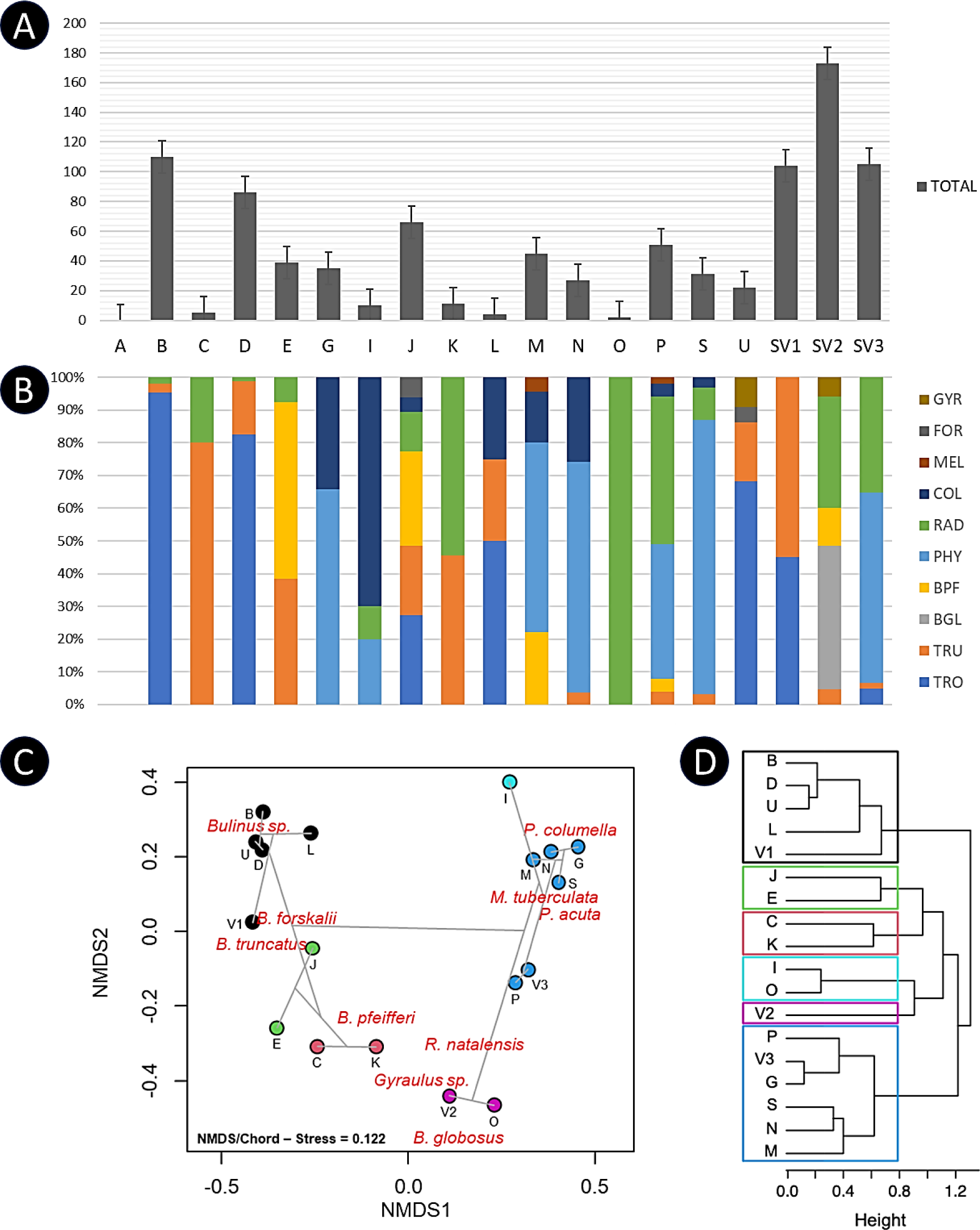
Visual representation of snail variables encompassing abundance, diversity and association with various sites. (A) Bar chart representing the proportional abundance for all snail species per site. (B) Snail diversity is shown by various colors per site and their relative abundance are shown as proportions for the respective bar. Color codes are given in the legend for each abbreviated snail species where GYR = *Gyraulus* sp., FOR = *B. forskalii*, MEL = *M. tuberculata*, COL = *P. columella*, RAD = *R. natalensis*, PHY = *P. acuta*, BPF = *B. pfeifferi*, BGL = *B. globosus*, TRU = *B. truncatus* and TRO = *B. tropicus*. Note that there is no bar for site A since no snails were found at this site. Significant differences between land use types are indicated with an asterisk. (C) NMDS plot and (D) UPGMA clustering dendrogram based on snail community data. NMDS plot with scores (circles) representing the 18 sites. Site A was omitted as no snails were present. The colors of the circles refer to the cluster the site belongs to. Snail species are indicated in red. The clustering dendrogram is visualized on the NMDS plot in grey lines. The distance between different sites represents their relatedness based on the snail community. The distance between sites and variables (here snail species) indicates the relatedness of the sites to the species.

No significant difference in snail abundance was found among the three conservation areas (Chiwawe, Middle section, and Lodge section), although there was high variation among sites within this region. Fig. 3A depicts the proportion of each snail species across the study area, highlighting significant diversity variations among different parts (ANOVA: *F*_4, 14_ = 3.18, *p* = 0.047). The Middle section and villages exhibited the highest diversity, while Chiwawe and the farm section had the lowest.

Concerning distribution, the NMDS analysis depicted a separation of sites along the horizontal axis, and the clustering dendrogram divided into two major branches (Fig. 3). One main group of sites was linked to *Bulinus* spp. and *B. pfeifferi*, while another main group consisted of sites characterized by *R. natalensis*, *M. tuberculata* and invasive species *P. columella* and *P. acuta*. Additionally, the NMDS plot revealed that sites in the farm area were associated with invasive species, and large dams (Sites M, P, and S) were also strongly linked to invasive snails. Sites within the conservation area showed an association with *B. pfeifferi* and *Bulinus* spp. On the other hand, village sites exhibited no clear relationship with each other based on the snail community.

### 3.2 Snail variables (infection rate)

Through shedding experiments, six cercarial morphotypes were identified. These included *Schistosoma* cercariae from a single *B. truncatus* snail at site V2, a village dam. Longifurcate pharyngeate distome subtype cercariae were found in 43 snails, including *B. tropicus*, *B. truncatus*, *B. globosus*, and *B. pfeifferi*. Avian schistosomes were released from two *B. truncatus*. Echinostome-type cercariae were released from 13 snails, involving *B. tropicus* and *B. pfeifferi*. Xiphidiocercariae-type cercariae were recovered from 13 *B. tropicus* snails, while amphistome-type cercariae were released from five snails, including *B. tropicus*, *B. truncatus*, and *B. forskalii*. In total, cercariae were identified after release from 104 out of 926 snails (11.2%). Among the 547 schistosome-competent snails, 88 were producing cercariae (16.1%). *Bulinus tropicus* exhibited the highest overall infection rate (76 out of 263, 28.9%), followed by *B. forskalii* (1 out of 5, 20.0%). The infection rate was 16 out of 168 (9.5%) for *R. natalensis*, 9 out of 131 (6.9%) for *B. truncatus*, and 2 out of 72 (2.8%) for *B. pfeifferi*. No cercariae were recovered from *P. acuta*, *P. columella*, *M. tuberculata*, and *Gyraulus* sp. during shedding experiments.

Table 1 provides a summary of infection prevalence per site based on molecular infection assessment with PCR, revealing that 70.20% (351/500) out of the extracted snails were infected with trematodes, ranging between 8.33% and 100% across sites. The overall infection prevalence among schistosome-competent snails, determined through RD-PCR, was 74.7%. Snail species with the highest infection rates were *B. forskalii* (100%), *B. tropicus* (93.4%), and *B. truncatus* (82.5%). *Bulinus globosus* had an infection rate of 46.7%, and *B. pfeifferi* showed an infection rate of 37.1%. No significant differences in infection prevalence were observed between different parts of Imire. Notably, three economically relevant infections were identified. Two *B. globosus* snails in the village area were infected with *Schistosoma mattheei* (site VX) and *S. haematobium* (site VY), as revealed by multiplex RD PCR. Additionally, one snail in the conservation area was infected with *Fasciola* sp., later identified as *Fasciola gigantica* through COI barcoding.

### 3.3 Environmental variables

Fig. 4 illustrates the variation in environmental variables across sites, with results from the principal component analysis (PCA). The PCA accounted for 29.2% of the variance in physicochemical variables between sites, with PC1 contributing 23.1% and PC2 contributing 16.2%. The variables carrying the most weight for PC1 were total dissolved solids (TDS) (35%), total suspended solids (TSS) (34%), and turbidity (33%). PC2 was predominantly associated with calcium and magnesium (40%). Sites in the farm area (i.e., G, I, O, and N) appeared on the right side of the 0-axis of PC1, indicating an association with higher TDS and conductivity levels. Along PC1, sites (A, C, and U) left of the 0.0 axis are typically eutrophic with high nitrate (32%), high phytoplankton biomass (23%), high turbidity (33%) and high level of blue-green algae (cyanobacteria) (28%). Several variables showed high correlations with each other, and the correlation plot angles reflect these associations. Distances among objects in the biplot do not represent approximations of their Euclidean distances in multidimensional space. Variables with high contribution were retained for redundancy analysis (RDA) to explore any associations with snail variables or the lack thereof (see section 3.5)

**Fig. 4.**
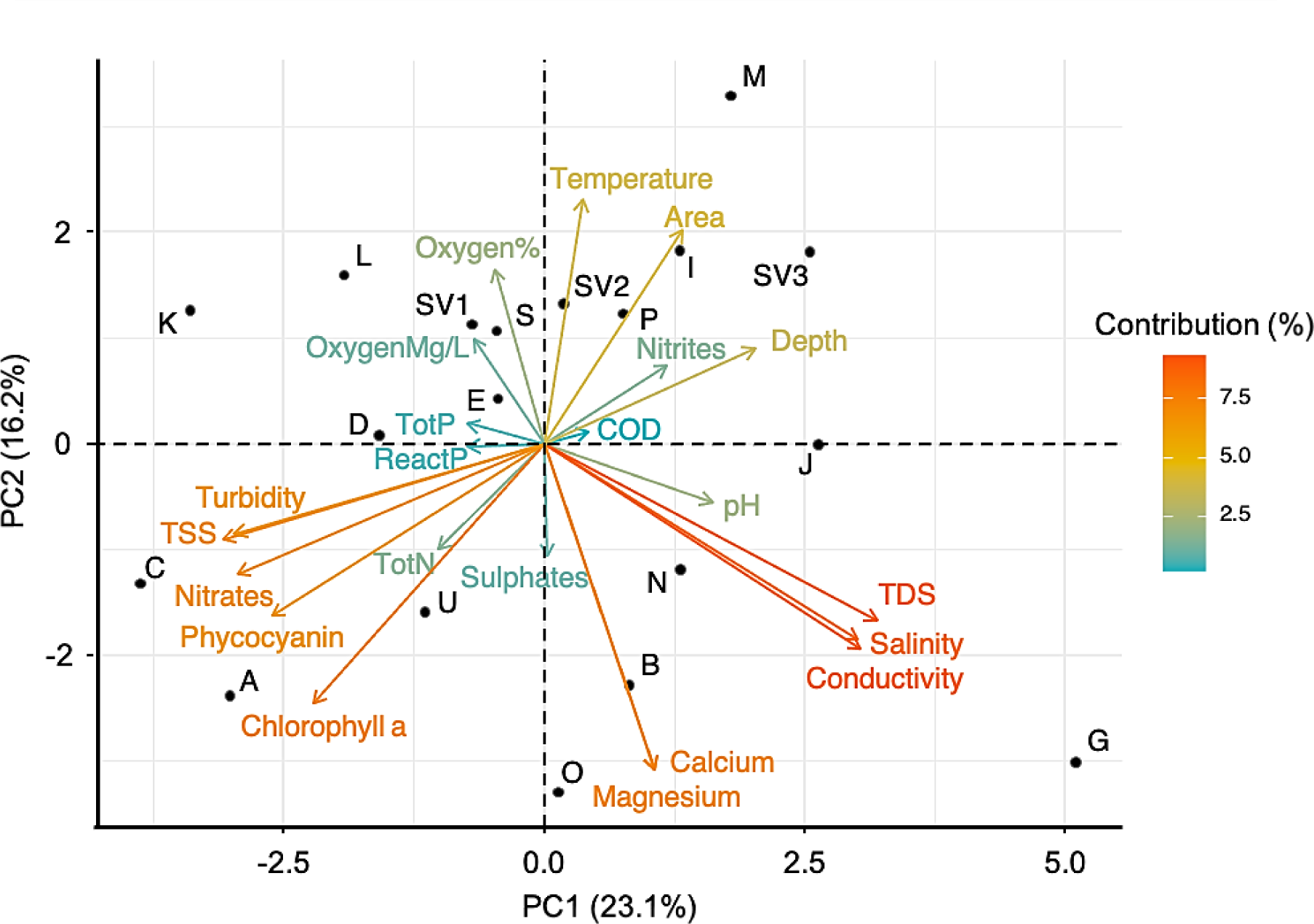
PCA correlation biplot (scaling = 2) visualizing the association among sites based on physical and chemical water variables. The scores (points) represent the 19 different sites, whereas the variables are shown as arrows. Contributions of the variables to the PCA are indicated by colors. Abbreviations stand for total dissolved solids (TDS), total suspended solids (TSS), total nitrogen (TotN), total phosphorus (TotP), reactive phosphorus (ReactP), and chemical oxygen demand (COD).

### 3.4 Macroinvertebrate and macrophyte communities

A total of 41 families of macroinvertebrates, with a diverse range of per-site counts (24 - 471 individuals), family richness (9 - 26), and Shannon diversity (1.148 - 2.586), were identified. The mean Shannon diversity was highest in Chiwawe and the Middle section, while it was lowest in the farm and lodge sections. However, the richness and diversity measurements between the different parts of the study area did not exhibit significant differences. Fig. 5 displays the NMDS and cluster analysis results. The NMDS plot does not reveal major groups based on the macroinvertebrate community, and the dispersion of sites across the plot does not appear to be linked to the different parts of the study area.

**Fig. 5.**
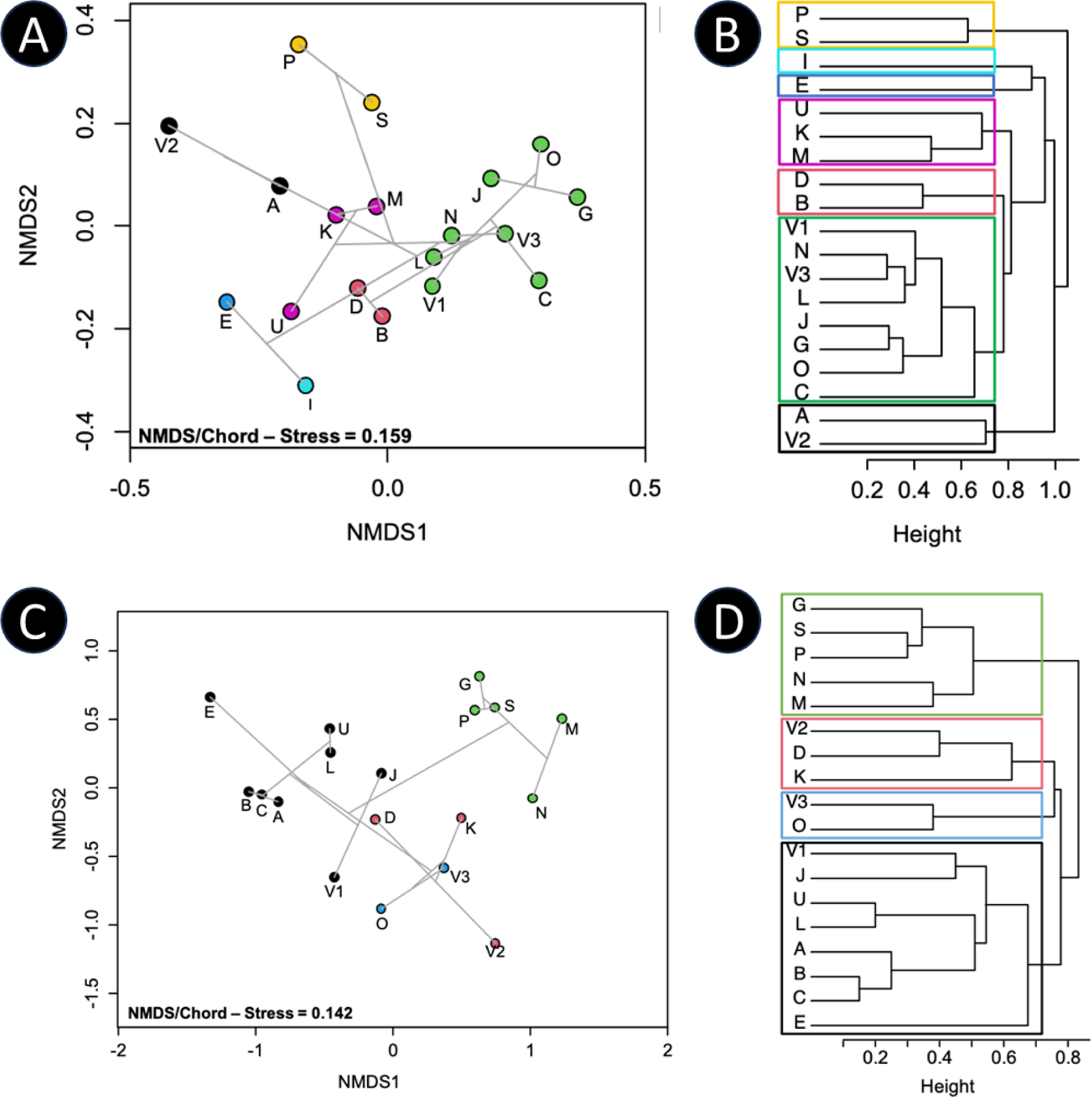
NMDS plots and UPGMA clustering dendrograms based on macroinvertebrate and macrophyte community data. (**A**) Macroinvertebrate NMDS plot with scores (circles) representing 19 sites, overlaid with (**B**) UPGMA clustering dendrogram (gray lines). Site I forms a distinct cluster due to its low macroinvertebrate count and richness. Elmidae is uniquely found at site E, forming a separate cluster. Other clusters exhibit similar family compositions and abundances: yellow (sites P and S), pink (sites K, M, and U), red (sites B and D), and green (sites with high Baetidae abundance). Sphaeriidae and Hydropsychidae are single occurrences only at site V2. (**C**) Macrophyte NMDS plot with scores (circles) representing 18 sites (Site I omitted). (**D**) UPGMA clustering dendrogram. Chiwawe sites (A, B, and C) cluster together, indicating shared macrophyte composition. The green cluster includes the farm, lodge, and upper middle section sites with *Lagarosiphon* sp. dominance. The red cluster comprises *Nymphaea caerulea* and *Stuckenia pectinata* abundance sites. The blue cluster shows *Typha capensis* prevalence and the black cluster contains Chiwawe and middle section sites with *Cladium mariscus* dominance.

A total of 21 macrophyte species were identified, and species richness and relative abundance were used to calculate Shannon diversity, while macrophyte cover was employed to estimate the abundance and cover of aquatic plants. Species richness varied between 3 and 9, and Shannon diversity ranged from 0.86 to 1.81. Submerged vegetation exhibited a cover range of 0 to 90%, with a mean of 30 ± 30%. Emergent vegetation ranged between 10 and 60%, with an average cover of 31 ± 15%, and floating vegetation cover varied from 0 to 70%, with a mean cover of 15 ± 16%. On the NMDS plot, sites exhibit a discernible separation along the horizontal axis (Fig. 5). The two significant aquatic plant species structuring the sites: *Lagarosiphon* sp. and *C. mariscus* (Fig. 5a) were retained for redundancy analysis (RDA) to explore any associations with snail variables (see subsection 3.5.3).

### 3.5 Snail variables and significant association or lack thereof

#### 3.5.1 Land use vs snail variables

Snail diversity and abundance were highest in the villages and lowest in the farm sites, although this was only significant for snail abundance (Χ^2^_(2,*N*=19)_= 15.44, p < 0.001). Tukey post hoc tests revealed significant differences between villages and farm sites and between villages and conservation sites. Land use did not significantly influence the infection rate, even though infection rates were also lowest in the farm sites as described above.

#### 3.5.2 Abiotic factors vs snail variables

To avoid cross-correlation, we selected TDS, calcium, chlorophyll A, phycocyanin, nitrate, algae, and turbidity—variables with the most weight in the PCA (Fig. 4)—for RDA. However, the resulting RDA did not show significance, suggesting that abiotic variables did not significantly explain the variation in the snail community. Given the known importance of abiotic factors in snail growth, all variables were included in a correlation plot employing forward selection. Two predictors emerged from the forward variable selection: depth (RDA1 = −0.97) and phycocyanin (RDA1 = 0.57, RDA2 = 0.82) (Fig. 6A). *Bulinus tropicus* and *B. truncatus* correlated with shallow depths, while *P. acuta* correlated with deeper depths. Additionally, *B. truncatus* showed a correlation with high cyanobacteria levels.

**Fig. 6.**
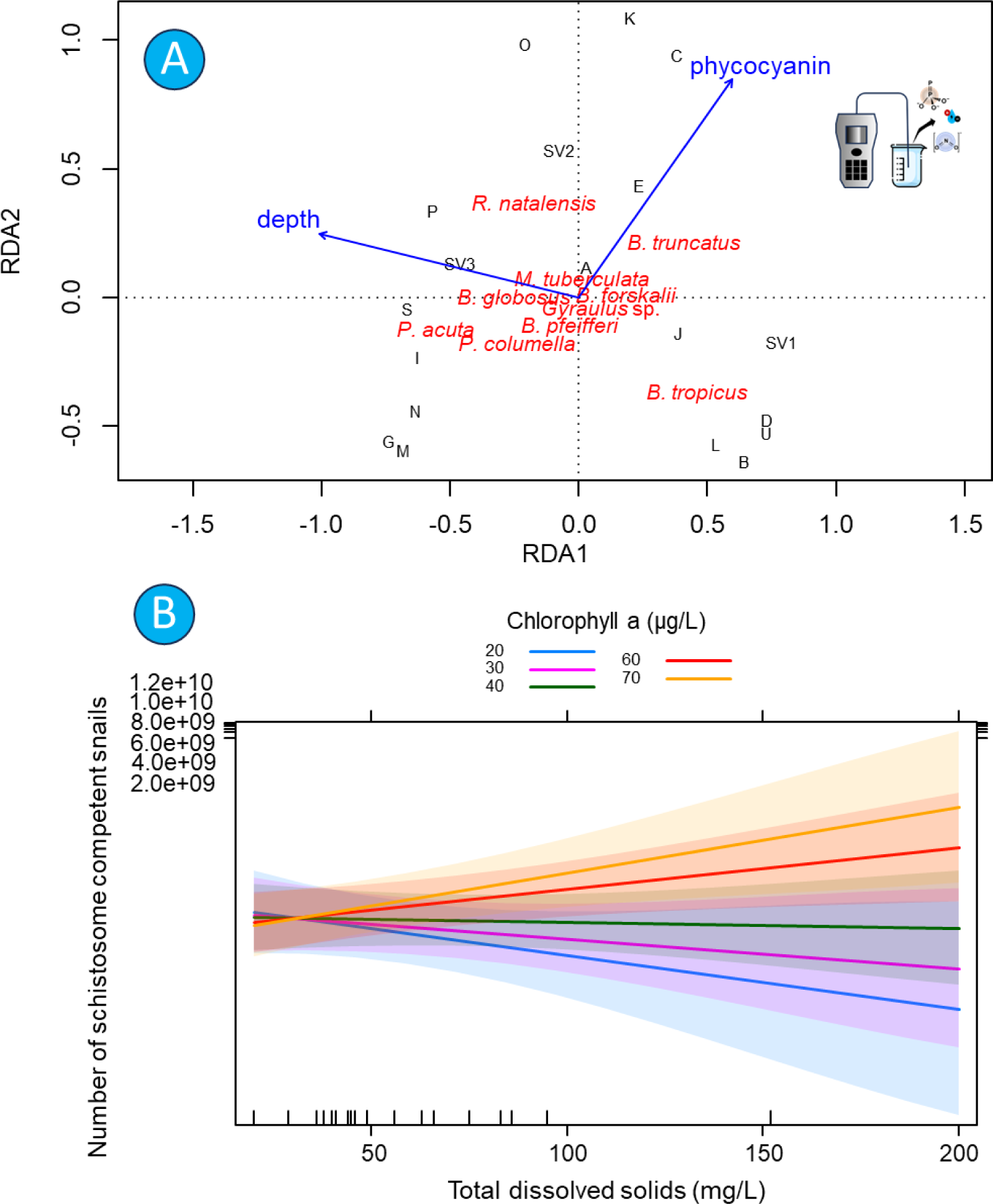
Visual representation of associations between some snail variables and water (abiotic) variables. (**A**) Correlation plot (Radj=0.277, p=0.001) for all abiotic variables resulting in only two predictors selected through the forward variable selection: depth (RDA1 = −0.97) and phycocyanin (RDA1 = 0.57, RDA2=0.82). Species scores on the plot were as follows: *P. acuta* (RDA1 = −0.56), *B. tropicus* (RDA1= 0.45, RDA2= 0.38), and *B. truncatus* (RDA1= 0.40). (**B**) Effect plot showing the association between the interaction of total dissolved solids (TDS) and chlorophyll A and the number of schistosome host snails. Colors indicate different chlorophyll A levels.

Again, all variables were used in linear mixed models, with only one model proving significant. This model revealed a significant association between the abundance of schistosome-competent snails and the interaction effect of TDS and chlorophyll A (Χ^2^_(1,*N*=19)_= 5.35, p = 0.02) (Fig. 6B). Specifically, at low chlorophyll A levels, TDS negatively impacted the abundance of schistosome host snails, while at high chlorophyll A levels, TDS had an increasing effect on their abundance. All other models were insignificant, indicating no significant associations between environmental variables and snail diversity, abundance, or infection rate.

#### 3.5.3 Macroinvertebrates vs snail variables

Macroinvertebrate diversity did not exhibit a statistically significant impact on the overall species richness, diversity, and infection rate of all snails. However, the Shannon diversity of macroinvertebrates demonstrated a significant positive correlation with the richness of schistosome host snails (Χ^2^_(1,*N*=19)_= 79.29, p < 0.001, Fig. 7A). Furthermore, the abundance of schistosome host snails displayed a significant positive association with the richness of macroinvertebrate families (Χ^2^_(1,*N*=19)_= 5.86, p = 0.016, Fig. 7B).

**Fig. 7.**
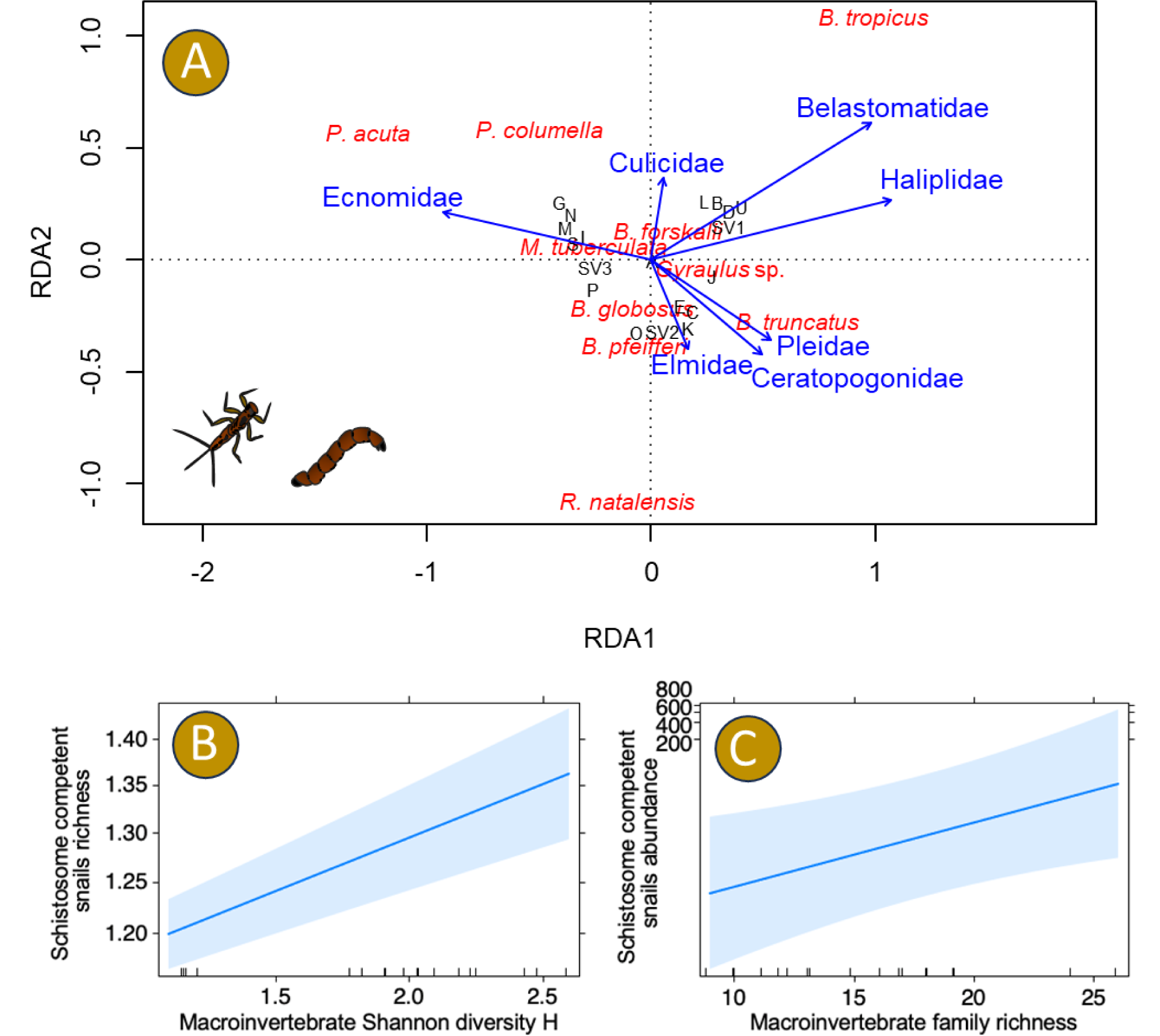
Visual representation of associations between some snail variables and macroinvertebrates. (**A**) Distance triplot of macroinvertebrates and snails using forward selection for macroinvertebrates groups. The triplot was significant (ANOVA: *F*7, 11 = 4.13, *p* = 0.001) and so were both RDA1 (ANOVA: *F*1, 11 = 13.86, *p* = 0.001) and RDA2 (ANOVA: *F*1, 11 = 9.27, *p* = 0.001) with Radj = 0.55. (**B**) Effect plots showing the significant (p < 0.001) association between macroinvertebrates Shannon diversity and the richness of schistosome host snails. (B) Effect plots showing the significant (p = 0.016) association between macroinvertebrate family richness and the abundance of schistosome host snails.

#### 3.5.4 Macrophyte vs snail variables

Following their perceived role in structuring the sites (see Fig 5B above), *Lagarosiphon major* and *C. mariscus* were therefore tested for association with snail variables using RDA. The resulting RDA was significant (ANOVA, F2,14 = 2.18; p = 0.031) and explained 23.7% of the variance, with an adjusted R2 value of 0.13 (Fig. 6b). The graph indicates that *B*. *tropicus* was indeed associated with the occurrence of *Cladium mariscus*, while *P. columella* and *P. acuta* were both most associated with *Lagarosiphon* sp. On the grand scale, macrophyte diversity did not show significant associations with the overall species richness, diversity, and infection rate of all snails as tested with LMMs. Macrophyte cover did not affect the infection rate either.

However, there were significant effects of macrophyte cover on snail counts and richness. Emergent vegetation cover significantly increased the richness of schistosome host snails ((Χ^2^_(1,*N*=19)_= 376.34, p < 0.001), Fig. 8B). Following the observations in previous multivariate analyses, it was additionally tested whether submerged vegetation (in our case mostly oxygen weed) was associated with *P. acuta* and lymnaeid snails *R. natalensis* and *P. columella*. Submerged vegetation cover significantly increased the richness (Χ^2^_(1,*N*=19)_= 4.01, p = 0.045), and abundance (Χ^2^_(1,*N*=19)_= 6.38, p = 0.012) of these snails (Figure 8C&D). Additional Pearson correlation tests were done revealing a significant correlation between snail abundance and floating vegetation cover (R = 0.6, p = 0.007, Fig. 8E). However, this correlation is skewed towards one outlier site (V2) which when removed from the analysis, no correlation exists.

**Fig. 8.**
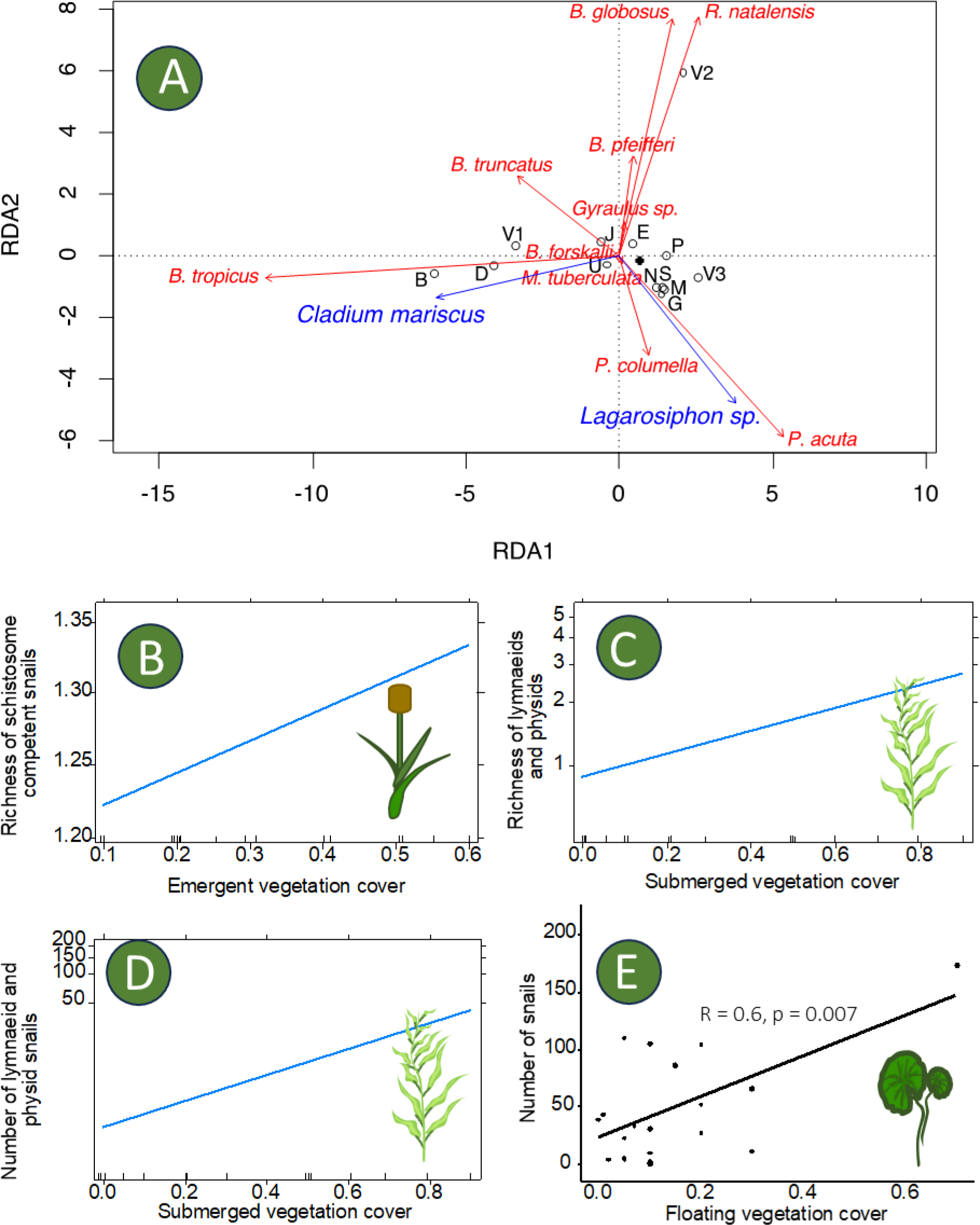
Visual representation of associations between some snail variables and water (abiotic) variables. (**A**) Distance triplot (scaling = 1) of the redundancy analysis. Sites (scores: black circles) are plotted in relation to the snail community of each site (response variables: red arrows) and macrophyte variables *Cladium mariscus* and *Lagarosiphon* sp. (explanatory variables: blue arrows). Distances among scores approximate their Euclidean distances and angles between response and explanatory variables reflect their correlation. The black star indicates sites K, O, L, and C that were positioned too closely to be displayed clearly. Effect plots showing (**B**) the association between emergent vegetation cover and the richness of schistosome host snails, (**C**) the association between submerged vegetation and the richness of Lymnaeidae and P. acuta and (**D**) the association between submerged vegetation and the abundance of Lymnaeidae and P. acuta. (**E**) shows a correlation plot between the abundance of snails and floating vegetation. Pearson correlation coefficient and p-value are shown on the plot.

## 4. Discussion

To achieve a more comprehensive and sustainable strategy for controlling the transmission of snail-borne diseases, it is crucial to deepen our understanding of the ecological dynamics involving snails as potential carriers of trematode parasites. Therefore, our investigation focused on elucidating the interplay among snail diversity, abundance of potential intermediate hosts, infection rates, and the land use surrounding man-made tropical water bodies in the Wedza district of Zimbabwe, alongside their abiotic and biotic features. The subsequent paragraphs will extensively discuss the key findings concerning the association between snail abundance, species richness, diversity, and infection rate in relation to 1) land use, 2) abiotic habitat characteristics, and 3) biotic habitat characteristics.

### 4.1 Snail-Land Use Relations

Human-induced alterations in land use, along with associated habitat loss and changes in nutrient cycles, stand out as key drivers of biodiversity decline (Powers & Jetz, 2019). However, their impacts vary across different taxonomic and functional groups (Graham et al., 2019; Newbold et al., 2019). In this study, we collected 10 species: *Bulinus tropicus*, *Bulinus truncatus*, *Bulinus globosus*, *Bulinus forskalii*, *Biomphalaria pfeifferi*, *Radix natalensis*, *Gyraulus* sp., *Melanoides tuberculata*, and the exotic invasive species *Pseudosuccinea columella* and *Physella acuta*. These findings align with a previous study by Mudavanhu et al. (in press) conducted in the same water bodies. Most of these species are short-lived pulmonate snails, including planorbids (e.g., *Bulinus* and *Biomphalaria*) and lymnaeids (i.e., *Radix*), which are ecological generalists capable of dispersing easily with or without hydrographic connections (Tabo et al., 2022).

Snail abundance varied significantly across the five parts of the study area characterized by different land uses, with the highest abundance observed in villages and the lowest in the farm section. Consistent with previous studies (Du et al., 2011; Lange et al., 2013; Oso & Odaibo, 2021), our findings underscore the profound impact of anthropogenic environmental changes on freshwater mollusc communities. Research suggests that snails tend to thrive in environments altered by human activities. For instance, in Lake Victoria, snail abundance was found to be greater at sites affected by human disturbance compared to undisturbed sites, indicating a preference for areas with human-water interactions (Lange et al., 2013). Additionally, schistosome host snails were found to be associated with disturbances linked to fishing villages (Lange et al., 2013). Both *Bulinus* and *Biomphalaria* have demonstrated a propensity for habitats affected by human disturbance, where they can exploit resources such as plastic waste for egg laying, proliferate in the absence of molluscivorous fish, or tolerate oxygen-depleted, polluted conditions (Stothard et al., 2017). This could explain the elevated abundance and richness of schistosome host snails observed in our sampled village dams (V1). Overall, snail diversity was highest in conserved areas and some village sites, while it was lowest in the farm section. Agriculture profoundly impacts stream morphology, water chemistry, siltation levels, nutrient dynamics, and hydrology (Paul & Meyer, 2001). It is also a significant driver of deforestation (Mereta et al., 2019; Paul & Meyer, 2001), leading to habitat loss/alteration and increased sunlight penetration, creating favorable conditions for snail survival. Both the village and farm sections in our study practice agriculture, but differences in intensity may have resulted in distinct outcomes. The farm section, primarily involved in commercial tobacco cultivation characterized by heavy fertilizer and pesticide use, exhibited low snail abundance and diversity. In contrast, traditional agriculture, the main livelihood source in the village sections, typically involves minimal inorganic agricultural inputs, leading to higher snail abundance and diversity.

### 4.2 Snail-Abiotic Relations

The intricate relationship between abiotic environmental conditions and the structure of snail communities has been a subject of extensive scientific investigation. Within our study, we observed variations in several water quality parameters across different habitat types. Specifically, three sites (A, C, and U) were identified as typically eutrophic, characterized by elevated levels of nitrate, phytoplankton biomass, turbidity, and cyanobacteria. Aquatic ecosystems exhibiting high nutrient and organic content, indicative of eutrophication, tend to be rich in trophic resources. This abundance may provide an advantage for generalist species of freshwater snails. The two exotic invasive species, *P. columella* and *P. acuta*, are known to be tolerant of eutrophication (Du et al., 2011; Grabner et al., 2014) and may thrive in these man-made reservoirs at the expense of native benthic fauna. However, the sensitivity of some species recorded in our study to eutrophication remains uncertain and could potentially have adverse effects. For instance, transcriptomic profiling has revealed potential molecular signals of eutrophication leading to developmental abnormalities in *Bellamya aeruginosa* (Lei et al., 2017). Sites within the commercial farm area showed an association with higher levels of total dissolved solids (TDS) and conductivity, potentially influencing the low species diversity observed, mainly dominated by *P. acuta*, a pond snail known for its resilience to high salinity. Numerous studies underscore the pivotal role of environmental conditions in shaping aquatic communities, a perspective that resonates with our findings. Through constrained ordinations and variable selection techniques, we unveiled significant correlations between snail community assemblages and both water depth and cyanobacteria levels. Water depth is recognized as a key determinant of general biodiversity and community structure among benthic invertebrates in aquatic ecosystems (Beisel et al., 1998), with several studies highlighting positive associations between shallow ponds and snails (Min et al., 2022).

In our investigation, *B. tropicus* and *B. truncatus* exhibited correlations with shallow depths, while *P. acuta* displayed an association with deeper depths. Previous research has emphasized the ecological importance of water depth in shaping the distribution of snail species (Olkeba et al., 2020; Oloyede et al., 2017). However, contrary to our findings, Olkeba et al., (2020) reported positive relationships between water depth and the occurrence of both *B. pfeifferi* and *B. globosus*, with a negative relationship observed for *Lymnaea truncatula*. Nevertheless, Mandahl-Barth, (1954) documented instances of *Biomphalaria smithi* and *Biomphalaria choanomphala* at depths of 4.3 m and 12.2 m, respectively, in Lake Edward and Lake Victoria, Uganda. Furthermore, in our study, *B. truncatus* exhibited a correlation with high cyanobacteria levels. This contrasts with the known negative impact of cyanobacterial phycocyanin on macroinvertebrate communities and its recognition as a promising candidate for microbial biopesticides targeting freshwater *Biomphalaria alexandrina* snails (Abd El-Ghany et al., 2018).

The interactive effects of total dissolved solids (TDS) and chlorophyll A revealed a nuanced relationship with the abundance of schistosome host snails. Specifically, we observed that at low chlorophyll A levels, increased TDS negatively impacted the abundance of these snails, whereas at high chlorophyll A levels, TDS had a stimulating effect on their abundance. Olkeba et al., (2020) similarly noted a negative correlation between chlorophyll-a content and the abundance of both *Biomphalaria sudanica* and *L. truncatula*. Chlorophyll-a serves as a crucial indicator of phytoplankton biomass in aquatic ecosystems, serving as a primary food source for snails. Salinity has also been identified as a significant factor influencing malacofauna, with higher TDS levels (above 17 gL^-1^) creating conditions unsuitable for mollusc survival (Sowa et al., 2019). Numerous studies conducted in saline ecosystems have demonstrated a direct relationship between macroinvertebrate species richness and salinization (e.g., Bäthe & Coring, 2011; Carver et al., 2009), generally indicating a decrease in biodiversity with increasing salinity. Braukmann & Böhme, (2011) observed that waters with the highest salt concentration exhibited extremely low species richness. Our findings suggest, to our knowledge for the first time, that chlorophyll-a, as a vital source of nourishment for snails, plays a critical role in conferring resilience to schistosome-competent snails against rising salinity levels.

### 4.3 Snail-Macroinvertebrate-Trematode Relations

We hypothesized that aquatic environments exhibiting a high diversity of macroinvertebrate communities and/or a substantial snail diversity would mitigate infection rates, following the widely acknowledged ‘decoy effect’. According to this concept, other organisms act as decoy hosts, diverting the parasite load away from focal hosts. Pre- and post-snail trematode larval stages, namely miracidia and cercariae, can infiltrate non-host snails, unsuitable final hosts, plants, and other macroinvertebrates where they cannot develop (Thieltges et al., 2008). Additionally, records exist for predation on these free-living parasite stages by oligochaete worms, dipteran larvae, and other insect larvae (Thieltges et al., 2008). Together, the decoy effect and predation contribute to the ‘dilution effect’, wherein high species diversity may decrease disease risk (Keesing et al., 2006; Keesing & Ostfeld, 2021). Contrary to our expectations, we found no discernible association between macroinvertebrate and/or snail diversity and infection rates.

Recent studies have scrutinized the universality of the dilution effect, positing that it only occurs under certain circumstances (Halliday & Rohr, 2019; Halsey & Miller, 2020; Strauss et al., 2015; Wood et al., 2017). The dilution effect is linked to the relationship between species richness and the relative densities of both focal host and non-host species (Strauss et al., 2015). Additionally, it is associated with the degree of nestedness in community assembly and the ecological characteristics of the parasites, such as their host specificity (Wood et al., 2014). Nonetheless, given the multi-host nature of certain trematodes, capable of infecting various unrelated snails, pinpointing which snails act as dilution hosts becomes challenging. To apply our results to the study of the dilution effect, it is necessary to assume that all surveyed reservoirs possess proportionally equal contamination with parasite eggs/miracidia, ensuring an equal infection pressure. However, this assumption is impractical since reservoirs, influenced by factors like location and size, are expected to face varying pressure from vertebrate hosts – considerations overlooked in our study. Clearly, our sampling design was not tailored to specifically address these nuances. Future investigations should attempt to quantify the infection pressure of a reservoir and integrate species abundance and functional roles of potential dilution organisms in addition to mere species diversity to unravel the dilution effect in trematode transmission. Species abundance determines encounter probability frequency, while the latter ensures the existence of diverse pathways for the dilution effect to manifest.

### 4.4 Do snails show a relation with macrophyte cover, diversity, or functional groups?

Aquatic plants play multifaceted roles in the ecology of snails. They serve as substrates for snail eggs, provide supplementary diets, and create suitable habitats by mitigating water velocity and tidal wave action (El Deeb et al., 2017; Plummer, 2005). High river flow can disrupt the availability of nutrients and substrates, thereby impacting snail growth and abundance, with different life cycle stages susceptible to being washed downstream and affecting overall population dynamics (Plummer, 2005). In our study, we identified a total of 21 different types of macrophytes across sites, categorized into submerged, emerged, and floating groups. These findings are consistent with previous research highlighting the importance of aquatic macrophytes in predicting freshwater snail species. For instance, our results revealed correlations between schistosome-competent snails and emergent vegetation cover, with redundancy analysis uncovering a specific association between *B. tropicus* and *Cladium mariscus*. Similar associations between schistosome-competent snails and emergent vegetation have been documented previously, such as *B. globosus* with *Cyperus* sp. and *Typha* sp., and *B. tropicus* with *Cyperus* sp. and *Polygonum* sp. (Chingwena et al., 2004; Pfukenyi et al., 2005). Other vegetation types have also been linked to these snails, including *B. globosus* and *B. pfeifferi* with submerged *Potamogeton* sp., and *B. tropicus* with floating *Nymphaea caerulea* (Chingwena et al., 2004; Pfukenyi et al., 2005). These findings suggest that associations between snail communities and local macrophytes may be specific to each ecosystem, emphasizing the importance of context-specific monitoring and management strategies tailored to individual reservoirs.

Furthermore, lymnaeids and physids showed associations with submerged vegetation. Specifically, *P. columella* and *P. acuta* were correlated with *Lagarosiphon major*. Despite being native to southern Africa, *L. major* (African curly-leaved waterweed, African elodea, oxygen weed) has become highly invasive worldwide, including in Europe (Stiers et al., 2011) and New Zealand (Riis et al., 2010). Its invasive success can be attributed to traits such as its vegetative growth habit, broad environmental tolerance, and high relative growth rate (Sutherland, 2004). The characteristic growth pattern of this plant leads to the formation of dense mats on and just below the water surface (DiTomaso et al., 2013), and as observed in our study. These mats can result in unintended fish kills due to oxygen depletion, hinder recreational activities such as boating, swimming, and angling, and obstruct hydroelectric and irrigation systems (DiTomaso et al., 2013; Matthews et al., 2012). Strong wave action can exacerbate the situation by dislodging weed mats, depositing rotting vegetation on beaches, ultimately reducing the ecosystem services provided by man-made reservoirs (DiTomaso et al., 2013). From a biodiversity perspective, the dense mats formed by *L. major* can block sunlight penetration to the lakebed, which is detrimental to aquatic invertebrates (DiTomaso et al., 2013). Interestingly, the two exotic invasive freshwater species, *P. columella* and *P. acuta*, appear to tolerate the adverse effects of this macrophyte. This exemplifies a “cascade of biological invasions” where the invasion success of an aquatic macrophyte promotes the abundance of exotic invasive freshwater snail species (Carolus et al., 2019), facilitated by damming activities that encourage the establishment of *L. major*. In their study, Carolus et al., (2019) reported a significant association between *P. columella* and *Eichhornia crassipes*, a floating water hyacinth species that also formed vast mats in the Lake Kariba – the world’s largest man-made reservoir volumetrically (Carolus et al., 2019).

## Conclusions

A surge in chemical control measures for schistosomiasis is imminent as nations strive to eliminate the disease by 2030, following China’s success as a benchmark, as reported by the World Health Organization. However, the widespread use of chemical control in rural waters, often intertwined with wildlife protected areas, can have extensive repercussions on aquatic ecosystem health and biodiversity.

To mitigate the reliance on chemical interventions for schistosomiasis control, our study proposes alternative strategies aimed at reducing snail populations. For instance, manipulating the abundance of emergent macrophytes, which have been found to correlate with schistosome-competent snails, holds promise in curtailing snail proliferation.

Moreover, our research highlights the vulnerability of these snails to high salinity levels in environments with low eutrophication. Thus, we advocate for minimizing nutrient inputs into reservoirs through improved agricultural and sewage disposal practices. This dual approach not only reduces schistosomiasis risk but also enhances ecosystem health and local biodiversity.

Furthermore, we identified that areas of shallow depth are conducive to hosting schistosome-competent hosts and suggest fencing off such areas to prevent animal and human contact with contaminated water. However, implementing this approach may pose challenges, particularly in regions where these sites serve as primary sources of drinking water for both animals and humans. Targeted chemical treatments at these sites could offer a temporary solution while allowing the reservoir to recover from the impacts of molluscicides.

## References

Abd El-Ghany, A. M., Salama, A., Abd El-Ghany, N. M., & Gharieb, R. M. A. (2018). New Approach for Controlling Snail Host of Schistosoma mansoni, Biomphalaria alexandrina with Cyanobacterial Strains-Derived C-Phycocyanin. *Vector Borne and Zoonotic Diseases (Larchmont*, N.Y.), 18(9), 464–468. 10.1089/vbz.2018.2274

Appleton, C., & Miranda, N. (2015). Two Asian Freshwater Snails Newly Introduced into South Africa and an Analysis of Alien Species Reported to Date. African Invertebrates, 56(1), 1–17. 10.5733/afin.056.0102

Bäthe, J., & Coring, E. (2011). Biological effects of anthropogenic salt-load on the aquatic Fauna: A synthesis of 17 years of biological survey on the rivers Werra and Weser. Limnologica, 41(2), 125–133. 10.1016/j.limno.2010.07.005

Becker, J. M., Ganatra, A. A., Kandie, F., Mühlbauer, L., Ahlheim, J., Brack, W., Torto, B., Agola, E. L., McOdimba, F., Hollert, H., Fillinger, U., & Liess, M. (2020). Pesticide pollution in freshwater paves the way for schistosomiasis transmission. Scientific Reports, 10(1), Article 1. 10.1038/s41598-020-60654-7

Beisel, J.-N., Usseglio-Polatera, P., Thomas, S., & Moreteau, J.-C. (1998). Stream community structure in relation to spatial variation: The influence of mesohabitat characteristics. Hydrobiologia, 389(1), 73–88. 10.1023/A:1003519429979

Braukmann, U., & Böhme, D. (2011). Salt pollution of the middle and lower sections of the river Werra (Germany) and its impact on benthic macroinvertebrates. Limnologica, 41(2), 113–124. 10.1016/j.limno.2010.09.003

Brown, D. S. (1994). Freshwater Snails of Africa and Their Medical Importance (2nd ed., Vol. 75). Taylor & Francis Ltd. 10.1016/0035-9203(81)90097-3

Carolus, H., Muzarabani, K. C., Hammoud, C., Schols, R., Volckaert, F. A. M., Barson, M., & Huyse, T. (2019). A cascade of biological invasions and parasite spillback in man-made Lake Kariba. Science of The Total Environment, 659, 1283–1292. 10.1016/j.scitotenv.2018.12.307

Carver, S., Storey, A., Spafford, H., Lynas, J., Chandler, L., & Weinstein, P. (2009). Salinity as a driver of aquatic invertebrate colonisation behaviour and distribution in the wheatbelt of Western Australia. Hydrobiologia, 617(1), 75–90. 10.1007/s10750-008-9527-5

Chingwena, G., Mukaratirwa, S., Chimbari, M., Kristensen, T. K., & Madsen, H. (2004). Population dynamics and ecology of freshwater gastropods in the highveld and lowveld regions of Zimbabwe, with emphasis on schistosome and amphistome intermediate hosts. African Zoology, 39(1), 55–62. 10.1080/15627020.2004.11407286

Civitello, D. J., Cohen, J., Fatima, H., Halstead, N. T., Liriano, J., McMahon, T. A., Ortega, C. N., Sauer, E. L., Sehgal, T., Young, S., & Rohr, J. R. (2015). Biodiversity inhibits parasites: Broad evidence for the dilution effect. Proceedings of the National Academy of Sciences of the United States of America, 112(28), 8667–8671. 10.1073/pnas.1506279112

Dai, J., Wang, W., Liang, Y., Li, H., Guan, X., & Zhu, Y. (2008). A novel molluscicidal formulation of niclosamide. Parasitology Research, 103(2), 405–412. 10.1007/s00436-008-0988-2

Day, J. A., Harrison, A. D., & de Moor, I. J. (2003). Guides to the Freshwater Invertebrates of Southern Africa: Insecta (Diptera) (Water Research Commission., Vol. 9).

de Moor, I. J., Day, J. A., & de Moor, F. C. (2003a). Guides to the Freshwater Invertebrates of Southern Africa: Insecta I (Ephemeroptera, Odonata&Plecoptera) (Water Research Commission., Vol. 7).

de Moor, I. J., Day, J. A., & de Moor, F. C. (2003b). Guides to the Freshwater Invertebrates of Southern Africa: Insecta II (Hemiptera, Megaloptera, Neuroptera, Trichoptera& Lepidoptera) (Water Research Commission., Vol. 8).

DiTomaso, J. M., Kyser, G. B., Oneto, S. R., Wilson, R. G., Orloff, S. B., Anderson, L. W., Wright, S. D., Roncoroni, J. A., Miller, T. L., & Prather, T. S. (2013). Weed Control in Natural Areas in the Western United States. University of California Weed Research and Information Center. https://books.google.be/books?id=QUC-mAEACAAJ

Du, L.-N., Li, Y., Chen, X.-Y., & Yang, J.-X. (2011). Effect of eutrophication on molluscan community composition in the Lake Dianchi (China, Yunnan). Limnologica, 41(3), 213–219. 10.1016/j.limno.2010.09.006

El Deeb, F., El-Shenawy, N., Soliman, M., & Mansour, S. (2017). Freshwater Snail Distribution Related to Physicochemical Parameters and Aquatic Macrophytes in Giza and Kafr El-Shiekh Governorates, Egypt. International Journal of Veterinary Science and Research, 3(1), 008–013. 10.17352/ijvsr.000015

Ellwanger, J. H., Kulmann-Leal, B., Kaminski, V. L., Valverde-Villegas, J. M., Veiga, A. B. G. D., Spilki, F. R., Fearnside, P. M., Caesar, L., Giatti, L. L., Wallau, G. L., Almeida, S. E. M., Borba, M. R., Hora, V. P. D., & Chies, J. A. B. (2020). Beyond diversity loss and climate change: Impacts of Amazon deforestation on infectious diseases and public health. Anais Da Academia Brasileira De Ciencias, 92(1), e20191375. 10.1590/0001-3765202020191375

Frandsen, F., & Christensen, N. O. (1984). An introductory guide to the identification of cercariae from African freshwater snails with special reference to cercariae of trematode species of medical and veterinary importance. Acta Tropica, 41(2), 181– 202.

Früh, D., Haase, P., & Stoll, S. (2017). Temperature drives asymmetric competition between alien and indigenous freshwater snail species, Physa acuta and Physa fontinalis. Aquatic Sciences, 79(1), 187–195. 10.1007/s00027-016-0489-9

Gerber, A., Cilliers, C. J., van Ginkel, C., & Glen, R. (2004). Easy identification of aquatic plants: A guide for the identification of water plants in and around South African impoundments. Department of Water Affairs.

Grabner, D. S., Mohamed, F. A. M. M., Nachev, M., Méabed, E. M. H., Sabry, A. H. A., & Sures, B. (2014). Invasion biology meets parasitology: A case study of parasite spill-back with Egyptian Fasciola gigantica in the invasive snail Pseudosuccinea columella. PloS One, 9(2), e88537. 10.1371/journal.pone.0088537

Graham, S. I., Kinnaird, M. F., O’Brien, T. G., Vågen, T.-G., Winowiecki, L. A., Young, T. P., & Young, H. S. (2019). Effects of land-use change on community diversity and composition are highly variable among functional groups. Ecological Applications, 29(7), e01973. 10.1002/eap.1973

Gyapong, J., & Boatin, B. (Eds.). (2016). Neglected Tropical Diseases—Sub-Saharan Africa. Springer International Publishing. 10.1007/978-3-319-25471-5

Gyasi, S. F., Boateng, A. A., Awuah, E., & Antwi, E. O. (2019). Ellucidating the incidence and the prevalence of Schistosomiasis spp infection in riparian communities of the Bui dam. Journal of Parasitic Diseases: Official Organ of the Indian Society for Parasitology, 43(2), 276–288. 10.1007/s12639-019-01089-4

Halliday, F. W., & Rohr, J. R. (2019). Measuring the shape of the biodiversity-disease relationship across systems reveals new findings and key gaps. Nature Communications, 10, 5032. 10.1038/s41467-019-13049-w

Halsey, S. J., & Miller, J. R. (2020). Maintenance of Borrelia burgdorferi among vertebrate hosts: A test of dilution effect mechanisms. Ecosphere, 11(2), e03048.

Halstead, N. T., Hoover, C. M., Arakala, A., Civitello, D. J., De Leo, G. A., Gambhir, M., Johnson, S. A., Jouanard, N., Loerns, K. A., McMahon, T. A., Ndione, R. A., Nguyen, K., Raffel, T. R., Remais, J. V., Riveau, G., Sokolow, S. H., & Rohr, J. R. (2018). Agrochemicals increase risk of human schistosomiasis by supporting higher densities of intermediate hosts. Nature Communications, 9(1), Article 1. 10.1038/s41467-018-03189-w

Hopkins, S. R., Wyderko, J. A., Sheehy, R. R., Belden, L. K., & Wojdak, J. M. (2013). Parasite predators exhibit a rapid numerical response to increased parasite abundance and reduce transmission to hosts. Ecology and Evolution, 3(13), 4427–4438. 10.1002/ece3.634

Hotez, P. J., Alvarado, M., Basáñez, M.-G., Bolliger, I., Bourne, R., Boussinesq, M., Brooker, S. J., Brown, A. S., Buckle, G., Budke, C. M., Carabin, H., Coffeng, L. E., Fèvre, E. M., Fürst, T., Halasa, Y. A., Jasrasaria, R., Johns, N. E., Keiser, J., King, C. H., … Naghavi, M. (2014). The global burden of disease study 2010: Interpretation and implications for the neglected tropical diseases. PLoS Neglected Tropical Diseases, 8(7), e2865. 10.1371/journal.pntd.0002865

Johansen, I. C., Moran, E. F., & Ferreira, M. U. (2023). The impact of hydropower dam construction on malaria incidence: Space-time analysis in the Brazilian Amazon. PLOS Global Public Health, 3(3), e0001683. 10.1371/journal.pgph.0001683

Johnson, P. T. J., Lund, P. J., Hartson, R. B., & Yoshino, T. P. (2009). Community diversity reduces Schistosoma mansoni transmission, host pathology and human infection risk. Proceedings of the Royal Society B: Biological Sciences, 276(1662), 1657–1663. 10.1098/rspb.2008.1718

Kalinda, C., Chimbari, M. J., & Mukaratirwa, S. (2017). Effect of temperature on the Bulinus globosus—Schistosoma haematobium system. Infectious Diseases of Poverty, 6(1), 57. 10.1186/s40249-017-0260-z

Kassambara, A., & Mundt, F. (2020). factoextra: Extract and Visualize the Results of Multivariate Data Analyses (1.0.7) [Computer software]. https://cran.r-project.org/web/packages/factoextra/index.html

Keesing, F., Holt, R. D., & Ostfeld, R. S. (2006). Effects of species diversity on disease risk. Ecology Letters, 9(4), 485–498.

Keesing, F., & Ostfeld, R. S. (2021). Dilution effects in disease ecology. Ecology Letters, 24(11), 2490–2505. 10.1111/ele.13875

Keighley, N., Ramwell, C., Sinclair, C., & Werner, D. (2021). Highly variable soil dissipation of metaldehyde can explain its environmental persistence and mobility. Chemosphere, 283, 131165. 10.1016/j.chemosphere.2021.131165

Kibret, S. (2018). Time to revisit how dams are affecting malaria transmission. The Lancet Planetary Health, 2(9), e378–e379. 10.1016/S2542-5196(18)30184-0

King, C. H., Sutherland, L. J., & Bertsch, D. (2015). Systematic Review and Meta-analysis of the Impact of Chemical-Based Mollusciciding for Control of Schistosoma mansoni and S. haematobium Transmission. PLOS Neglected Tropical Diseases, 9(12), e0004290. 10.1371/journal.pntd.0004290

Kokaliaris, C., Garba, A., Matuska, M., Bronzan, R. N., Colley, D. G., Dorkenoo, A. M., Ekpo, U. F., Fleming, F. M., French, M. D., Kabore, A., Mbonigaba, J. B., Midzi, N., Mwinzi, P. N. M., N’Goran, E. K., Polo, M. R., Sacko, M., Tchuenté, L.-A. T., Tukahebwa, E. M., Uvon, P. A., … Vounatsou, P. (2022). Effect of preventive chemotherapy with praziquantel on schistosomiasis among school-aged children in sub-Saharan Africa: A spatiotemporal modelling study. The Lancet Infectious Diseases, 22(1), 136–149. 10.1016/S1473-3099(21)00090-6

Lai, Y.-S., Biedermann, P., Ekpo, U. F., Garba, A., Mathieu, E., Midzi, N., Mwinzi, P., N’Goran, E. K., Raso, G., Assaré, R. K., Sacko, M., Schur, N., Talla, I., Tchuenté, L.-A. T., Touré, S., Winkler, M. S., Utzinger, J., & Vounatsou, P. (2015). Spatial distribution of schistosomiasis and treatment needs in sub-Saharan Africa: A systematic review and geostatistical analysis. The Lancet. Infectious Diseases, 15(8), 927–940. 10.1016/S1473-3099(15)00066-3

Lange, C., Kristensen, T., & Madsen, H. (2013). Gastropod diversity, distribution and abundance in habitats with and without anthropogenic disturbances in Lake Victoria, Kenya. African Journal of Aquatic Science, 38(3), 295–304. 10.2989/16085914.2013.797380

Larson, M. D., & Ross Black, A. (2016). Assessing interactions among native snails and the invasive New Zealand mud snail, Potamopyrgus antipodarum, using grazing experiments and stable isotope analysis. Hydrobiologia, 763(1), 147–159. 10.1007/s10750-015-2369-z

Lawler, O. K., Allan, H. L., Baxter, P. W. J., Castagnino, R., Tor, M. C., Dann, L. E., Hungerford, J., Karmacharya, D., Lloyd, T. J., López-Jara, M. J., Massie, G. N., Novera, J., Rogers, A. M., & Kark, S. (2021). The COVID-19 pandemic is intricately linked to biodiversity loss and ecosystem health. The Lancet Planetary Health, 5(11), e840–e850. 10.1016/S2542-5196(21)00258-8

Lei, K., Qiao, F., Liu, Q., Wei, Z., An, L., Qi, H., Cui, S., & LeBlanc, G. A. (2017). Preliminary evidence for snail deformation from a Eutrophic lake. Environmental Toxicology and Pharmacology, 53, 219–226. 10.1016/j.etap.2017.06.019

Malan, H. L., Appleton, C. C., Day, J. A., & Dini, J. (2009). Review: Wetlands and invertebrate disease hosts: Are we asking for trouble? Water SA, 35(5), Article 5. 10.4314/wsa.v35i5.49202

Maldonado, M. A., & Martín, P. R. (2019). Dealing with a hyper-successful neighbor: Effects of the invasive apple snail Pomacea canaliculata on exotic and native snails in South America. Current Zoology, 65(3), 225–235. 10.1093/cz/zoy060

Mandahl-Barth, G. (1954). The freshwater mollusks of Uganda and adjacent territories. Koninklijk Museum van Belgisch-Congo.

Mandahl-Barth, G. (1962). Key to the identification of east and central African freshwater snails of medical and veterinary importance. Bulletin of the World Health Organization, 27(1), 135–150.

Matthews, J., Beringen, R., Collas, F. P. L., Koopman, K. R., Odé, B., Pot, R., Sparrius, L. B., & Verbrugge, L. N. H. (2012). Knowledge document for risk analysis of the non-native Curly Waterweed (Lagarosiphon major) in the Netherlands. Reports Environmental Science, 414; Department of Environmental Science: Nijmegen, The Netherlands. https://repository.ubn.ru.nl/bitstream/handle/2066/103462/103462.pdf

Mereta, S. T., Bedewi, J., Yewhalaw, D., Mandefro, B., Abdie, Y., Tegegne, D., Birke, W., Mulat, W. L., & Kloos, H. (2019). Environmental determinants of distribution of freshwater snails and trematode infection in the Omo Gibe River Basin, southwest Ethiopia. Infectious Diseases of Poverty, 8(1), 93. 10.1186/s40249-019-0604-y

Midzi, N., Mduluza, T., Chimbari, M. J., Tshuma, C., Charimari, L., Mhlanga, G., Manangazira, P., Munyati, S. M., Phiri, I., Mutambu, S. L., Midzi, S. S., Ncube, A., Muranzi, L. P., Rusakaniko, S., & Mutapi, F. (2014). Distribution of Schistosomiasis and Soil Transmitted Helminthiasis in Zimbabwe: Towards a National Plan of Action for Control and Elimination. PLoS Neglected Tropical Diseases, 8(8), e3014. 10.1371/journal.pntd.0003014

Min, F., Wang, J., Liu, X., Yuan, Y., Guo, Y., Zhu, K., Chai, Z., Zhang, Y., & Li, S. (2022). Environmental Factors Affecting Freshwater Snail Intermediate Hosts in Shenzhen and Adjacent Region, South China. Tropical Medicine and Infectious Disease, 7(12), 426. 10.3390/tropicalmed7120426

Newbold, T., Adams, G. L., Albaladejo Robles, G., Boakes, E. H., Braga Ferreira, G., Chapman, A. S. A., Etard, A., Gibb, R., Millard, J., Outhwaite, C. L., & Williams, J. J. (2019). Climate and land-use change homogenise terrestrial biodiversity, with consequences for ecosystem functioning and human well-being. Emerging Topics in Life Sciences, 3(2), 207–219. 10.1042/ETLS20180135

Oksanen, J., Blanchet, F. G., Friendly, M., Kindt, R., Legendre, P., McGlinn, D., Minchin, P., O’Hara, R., Simpson, G., Solymos, P., Stevens, M., Szöcs, E., & Wagner, H. (2022). Vegan: Community Ecology Package version 2.5-7.

Oldham, R. S., Keeble, J., & Jeffcote, M. J. S. S. A. M. (2000). Evaluating the suitability of habitat for the great crested newt (Triturus cristatus). Herpetological Journal, 10(4), 143–155.

Olkeba, B. K., Boets, P., Mereta, S. T., Yeshigeta, M., Akessa, G. M., Ambelu, A., & Goethals, P. L. M. (2020). Environmental and biotic factors affecting freshwater snail intermediate hosts in the Ethiopian Rift Valley region. Parasites & Vectors, 13(1), 292. 10.1186/s13071-020-04163-6

Oloyede, O. O., Otarigho, B., & Morenikeji, O. (2017). Diversity, distribution and abundance of freshwater snails in Eleyele dam, Ibadan, south-west Nigeria. Zoology and Ecology, 27(1), 35–43. 10.1080/21658005.2016.1245934

Oso, O. G., & Odaibo, A. B. (2021). Land use/land cover change, physico-chemical parameters and freshwater snails in Yewa North, Southwestern Nigeria. PLOS ONE, 16(2), e0246566. 10.1371/journal.pone.0246566

Paul, M. J., & Meyer, J. L. (2001). Streams in the Urban Landscape. Annual Review of Ecology and Systematics, 32(1), 333–365. 10.1146/annurev.ecolsys.32.081501.114040

Pfukenyi, D. M., Monrad, J., & Mukaratirwa, S. (2005). Epidemiology and control of trematode infections in cattle in Zimbabwe: A review. Journal of the South African Veterinary Association, 76(1), 9–17. 10.4102/jsava.v76i1.387

Pica-Mattoccia, L., & Cioli, D. (2004). Sex- and stage-related sensitivity of Schistosoma mansoni to in vivo and in vitro praziquantel treatment. International Journal for Parasitology, 34(4), 527–533. 10.1016/j.ijpara.2003.12.003

Plummer, M. L. (2005). Impact of Invasive Water Hyacinth (Eichhornia crassipes) on Snail Hosts of Schistosomiasis in Lake Victoria, East Africa. EcoHealth, 2(1), 81–86. 10.1007/s10393-004-0104-8

Powers, R. P., & Jetz, W. (2019). Global habitat loss and extinction risk of terrestrial vertebrates under future land-use-change scenarios. Nature Climate Change, 9(4), Article 4. 10.1038/s41558-019-0406-z

R Development Core Team. (2018). R: A language and environment for statistical computing. R Foundation for Statistical Computing, Vienna, Austria. https://www.r-project.org/

Riis, T., Lambertini, C., Olesen, B., Clayton, J. S., Brix, H., & Sorrell, B. K. (2010). Invasion strategies in clonal aquatic plants: Are phenotypic differences caused by phenotypic plasticity or local adaptation? Annals of Botany, 106(5), 813–822. 10.1093/aob/mcq176

Rollinson, D., Knopp, S., Levitz, S., Stothard, J. R., Tchuem Tchuenté, L.-A., Garba, A., Mohammed, K. A., Schur, N., Person, B., Colley, D. G., & Utzinger, J. (2013). Time to set the agenda for schistosomiasis elimination. Acta Tropica, 128(2), 423–440. 10.1016/j.actatropica.2012.04.013

Schols, R., Carolus, H., Hammoud, C., Mulero, S., Mudavanhu, A., & Huyse, T. (2019). A rapid diagnostic multiplex PCR approach for xenomonitoring of human and animal schistosomiasis in a “One Health” context. Transactions of the Royal Society of Tropical Medicine and Hygiene, 113(11), 722–729. 10.1093/trstmh/trz067

Sokolow, S. H., Jones, I. J., Jocque, M., La, D., Cords, O., Knight, A., Lund, A., Wood, C. L., Lafferty, K. D., Hoover, C. M., Collender, P. A., Remais, J. V., Lopez-Carr, D., Fisk, J., Kuris, A. M., & De Leo, G. A. (2017). Nearly 400 million people are at higher risk of schistosomiasis because dams block the migration of snail-eating river prawns. Philosophical Transactions of the Royal Society of London. Series B, Biological Sciences, 372(1722), 20160127. 10.1098/rstb.2016.0127

Sokolow, S. H., Wood, C. L., Jones, I. J., Lafferty, K. D., Kuris, A. M., Hsieh, M. H., & De Leo, G. A. (2018). To Reduce the Global Burden of Human Schistosomiasis, Use “Old Fashioned” Snail Control. Trends in Parasitology, 34(1), 23–40. 10.1016/j.pt.2017.10.002

Sokolow, S. H., Wood, C. L., Jones, I. J., Swartz, S. J., Lopez, M., Hsieh, M. H., Lafferty, K. D., Kuris, A. M., Rickards, C., & De Leo, G. A. (2016). Global Assessment of Schistosomiasis Control Over the Past Century Shows Targeting the Snail Intermediate Host Works Best. PLoS Neglected Tropical Diseases, 10(7), e0004794. 10.1371/journal.pntd.0004794

Sowa, A., Krodkiewska, M., Halabowski, D., & Lewin, I. (2019). Response of the mollusc communities to environmental factors along an anthropogenic salinity gradient. The Science of Nature, 106(11), 60. 10.1007/s00114-019-1655-4

Stals, R., & de Moor, I. J. (2007). Guides to the Freshwater Invertebrates of Southern Africa: Insecta (Coleoptera) (Water Research Commission., Vol. 10).

Steinmann, P., Keiser, J., Bos, R., Tanner, M., & Utzinger, J. (2006). Schistosomiasis and water resources development: Systematic review, meta-analysis, and estimates of people at risk. The Lancet Infectious Diseases, 6(7), 411–425. 10.1016/S1473-3099(06)70521-7

Stiers, I., Njambuya, J., & Triest, L. (2011). Competitive abilities of invasive *Lagarosiphon major* and native *Ceratophyllum demersum* in monocultures and mixed cultures in relation to experimental sediment dredging. Aquatic Botany, 95(2), 161–166. 10.1016/j.aquabot.2011.05.011

Stothard, J. R., Campbell, S. J., Osei-Atweneboana, M. Y., Durant, T., Stanton, M. C., Biritwum, N.-K., Rollinson, D., Ombede, D. R. E., & Tchuem-Tchuenté, L.-A. (2017). Towards interruption of schistosomiasis transmission in sub-Saharan Africa: Developing an appropriate environmental surveillance framework to guide and to support ‘end game’ interventions. Infectious Diseases of Poverty, 6(1), 10. 10.1186/s40249-016-0215-9

Strauss, A. T., Civitello, D. J., Cáceres, C. E., & Hall, S. R. (2015). Success, failure and ambiguity of the dilution effect among competitors. Ecology Letters, 18(9), 916–926. 10.1111/ele.12468

Sutherland, S. (2004). What makes a weed a weed: Life history traits of native and exotic plants in the USA. Oecologia, 141(1), 24–39. 10.1007/s00442-004-1628-x

Tabo, Z., Neubauer, T. A., Tumwebaze, I., Stelbrink, B., Breuer, L., Hammoud, C., & Albrecht, C. (2022). Factors Controlling the Distribution of Intermediate Host Snails of Schistosoma in Crater Lakes in Uganda: A Machine Learning Approach. Frontiers in Environmental Science, 10. https://www.frontiersin.org/articles/10.3389/fenvs.2022.871735

Thieltges, D. W., Bordalo, M. D., Hernández, A. C., Prinz, K., & Jensen, K. T. (2008). Ambient fauna impairs parasite transmission in a marine parasite-host system. Parasitology, 135(9), 1111–1116. 10.1017/S0031182008004526

Toledo, R., & Fried, B. (2014). Digenetic trematodes. Springer.

Turner, A. M., Turner, R. R., & Ray, S. R. (2007). Competition and intraguild egg predation among freshwater snails: Re-examining the mechanism of interspecific interactions. Oikos, 116(11), 1895–1903. 10.1111/j.0030-1299.2007.15883.x

Webbe, G., & el Hak, S. (1990). Progress in the control of schistosomiasis in Egypt 1985-1988. Transactions of the Royal Society of Tropical Medicine and Hygiene, 84(3), 394–400. 10.1016/0035-9203(90)90334-b

Wenseleers, T. (2016). Export: R package for streamlined export of graphs and data tables. GitHub. https://github.com/tomwenseleers/export

WHO. (2020). Ending the neglect to attain the Sustainable Development Goals: A road map for neglected tropical diseases 2021–2030. World Health Organization. https://www.who.int/publications-detail-redirect/9789240010352

Wood, C. L., Lafferty, K. D., DeLeo, G., Young, H. S., Hudson, P. J., & Kuris, A. M. (2014). Does biodiversity protect humans against infectious disease? Ecology, 95(4), 817– 832. 10.1890/13-1041.1

Wood, C. L., McInturff, A., Young, H. S., Kim, D., & Lafferty, K. D. (2017). Human infectious disease burdens decrease with urbanization but not with biodiversity. Philosophical Transactions of the Royal Society B: Biological Sciences, 372(1722), 20160122. 10.1098/rstb.2016.0122

Yang, G.-J., Sun, L.-P., Hong, Q.-B., Zhu, H.-R., Yang, K., Gao, Q., & Zhou, X.-N. (2012). Optimizing molluscicide treatment strategies in different control stages of schistosomiasis in the People’s Republic of China. Parasites & Vectors, 5(1), 260. 10.1186/1756-3305-5-260

Yang, Q., Liu, M., Huang, D., Xiong, W., Yu, Q., Guo, T., & Wei, Q. (2020). A Survey and Ecological Risk Assessment of Niclosamide and Its Degradation Intermediatein Wucheng Waters within Poyang Lake Basin, China. Polish Journal of Environmental Studies, 30(1), 433–451. 10.15244/pjoes/117695

